# 15-keto-prostaglandin E_2_ activates host peroxisome proliferator-activated receptor gamma (PPAR-γ) to promote *Cryptococcus neoformans* growth during infection

**DOI:** 10.1101/113167

**Authors:** Robert J. Evans, Katherine Pline, Catherine A. Loynes, Sarah Needs, Maceler Aldrovandi, Jens Tiefenbach, Ewa Bielska, Rachel E. Rubino, Christopher J. Nicol, Robin C. May, Henry M. Krause, Valerie B. O’Donnell, Stephen A. Renshaw, Simon A. Johnston

**Affiliations:** Bateson Centre, Firth Court, University of Sheffield, S10 2TN, UK; Department of Infection, Immunity and Cardiovascular Disease, Medical School, University of Sheffield, S10 2RX, UK; Institute of Microbiology and Infection, School of Biosciences, University of Birmingham, Birmingham, B15 2TT, UK; Systems Immunity Research Institute, and Division of Infection and Immunity, School of Medicine, Cardiff University, Cardiff CF14 4XN, UK; Banting and Best Department of Medical Research, The Terrence Donnelly Centre for Cellular and Biomolecular Research (CCBR), University of Toronto, Toronto, Ontario, Canada, InDanio Bioscience Inc., Toronto, Ontario, Canada; Department of Pathology and Molecular Medicine, Queen’s University, Kingston, ON, Canada

**Author notes:** Author for correspondence: Simon A. Johnston.

## Abstract

*Cryptococcus neoformans* is one of the leading causes of invasive fungal infection in humans worldwide. *C. neoformans* uses macrophages as a proliferative niche to increase infective burden and avoid immune surveillance. However, the specific mechanisms by which *C. neoformans* manipulates host immunity to promote its growth during infection remain ill-defined. Here we demonstrate that eicosanoid lipid mediators manipulated and/or produced by *C. neoformans* play a key role in regulating pathogenesis. *C. neoformans* is known to secrete several eicosanoids that are highly similar to those found in vertebrate hosts. Using eicosanoid deficient cryptococcal mutants *Δplb1* and *Δlac1*, we demonstrate that prostaglandin E_2_ is required by *C. neoformans* for proliferation within macrophages and *in vivo* during infection. Genetic and pharmacological disruption of host PGE_2_ synthesis is not required for promotion of cryptococcal growth by eicosanoid production. We find that PGE_2_ must be dehydrogenated into 15-keto-PGE_2_ to promote fungal growth, a finding that implicated the host nuclear receptor PPAR-*γ*. *C. neoformans* infection of macrophages activates host PPAR-*γ* and its inhibition is sufficient to abrogate the effect of 15-keto-PGE_2_ in promoting fungal growth during infection. Thus, we describe the first mechanism of reliance on pathogen-derived eicosanoids in fungal pathogenesis and the specific role of 15-keto-PGE_2_ and host PPAR-*γ* in cryptococcosis.

**Author Summary:** *Cryptococcus neoformans* is an opportunistic fungal pathogen that is responsible for significant numbers of deaths in the immunocompromised population worldwide. Here we address whether eicosanoids produced by *C. neoformans* manipulate host innate immune cells during infection. *Cryptococcus neoformans* produces several eicosanoids that are notable for their similarity to vertebrate eicosanoids, it is therefore possible that fungal-derived eicosanoids may provoke physiological effects in the host. Using a combination of *in vitro* and *in vivo* infection models we identify a specific eicosanoid species - prostaglandin E_2_ – that is required by *C. neoformans* for growth during infection. We subsequently show that prostaglandin E_2_ must be converted to 15-keto-prostaglandin E_2_ within the host before it has these effects. Furthermore, we find that prostaglandin E_2_/15-keto-prostaglandin E_2_ mediated virulence is via activation of host PPAR-*γ* – an intracellular eicosanoid receptor known to interact with 15-keto-PGE_2_.

## Introduction

*Cryptococcus neoformans* is an opportunistic pathogen that infects individuals who have severe immunodeficiencies such as late-stage HIV AIDS. *C. neoformans* is estimated to infect 278,000 individuals each year resulting in 181,000 deaths (1,2). *C. neoformans* infection begins in the lungs where the fungus is phagocytosed by host macrophages. Macrophages must become activated by further inflammatory signals from the host immune system before they can effectively kill *C. neoformans* (3,4). When this does not occur *C. neoformans* proliferates rapidly intracellularly and may use the macrophage to disseminate to the central nervous system leading to fatal cryptococcal meningitis (5–9).

Eicosanoids are an important group of lipid inflammatory mediators produced by innate immune cells such as macrophages. Eicosanoids are a diverse group of potent signalling molecules that have a short range of action and signal through autocrine and paracrine routes. Macrophages produce large amounts of a particular group of eicosanoids called prostaglandins during microbial infection (10,11). Prostaglandins have a number of physiological effects throughout the body, but in the context of immunity they are known to strongly influence the inflammatory state (12). The prostaglandins PGE_2_ and PGD_2_ are the best-studied eicosanoid inflammatory mediators. During infection, macrophages produce both PGE_2_ and PGD_2_ to which, via autocrine routes, they are highly responsive (12). In vertebrate immunity, the synthesis of eicosanoids such as PGE_2_ is carefully regulated by feedback loops to ensure that the potent effects of these molecules are properly constrained. Exogenous sources of eicosanoids within the body, such as from eicosanoid-producing parasites (13) or tumours that overproduce eicosanoids (14,15), can disrupt host inflammatory signaling as they are not subject to the same regulation.

It is well known that *C. neoformans* produces its own eicosanoid species. These fungal-derived eicosanoids are indistinguishable from those produced by vertebrates (16–18). Only two *Cryptococcus* enzymes are known to be associated with cryptococcal eicosanoid synthesis - phospholipase B1 and laccase (18,19). Deletion of phospholipase B1 reduces secreted levels of all eicosanoids produced by *C. neoformans* suggesting that it has high level role in eicosanoid synthesis (19), perhaps fulfilling the role of phospholipase A_2_ in higher organisms. Deletion of laccase results in reduced levels of PGE_2_ but other eicosanoids are unaffected suggesting that laccase has putative PGE_2_ synthase activity (18). *C. neoformans* produces eicosanoids during infection, these eicosanoids are indistinguishable from host eicosanoids so it is possible that *C. neoformans* is able to manipulate the host inflammatory state during infection by directly manipulating host eicosanoid signaling.

It has previously been reported that the inhibition of prostaglandin E_2_ receptors EP2 and EP4 during murine pulmonary infection leads to better host survival accompanied by a shift towards Th1/M1 macrophage activation, however it was not determined if PGE_2_ was derived from the host or the fungus (20). Therefore, a key aspect of *C. neoformans* pathogenesis remains unanswered: do eicosanoids produced by *C. neoformans* manipulate host innate immune cells function during infection?

We have previously shown that the eicosanoid deficient strain *plb1* has reduced proliferation and survival within macrophages (21). We hypothesised that eicosanoids produced by *C. neoformans* support intracellular proliferation within macrophages and subsequently promote pathogenesis. To address this hypothesis, we combined *in vitro* macrophage infection assays with our previous published *in vivo* zebrafish model of cryptococcosis (22). We found that PGE_2_ was sufficient to promote growth of *Δplb1* and *Δlac1* independent of host PGE_2_ production, *in vitro* and *in vivo*. We show that the effects of PGE_2_ in cryptococcal infection are mediated by its dehydrogenated form, 15-keto-PGE_2_. Finally, we determine that 15-keto-PGE_2_ promotes *C. neoformans* infection via the activation of the host nuclear transcription factor PPAR-*γ*, demonstrating that 15-keto-PGE_2_ and PPAR-*γ* are new factors in cryptococcal infection

## Results

### Prostaglandin E_2_ is required for C. neoformans growth in macrophages

We have previously shown that the *C. neoformans* mutant strain *Δplb1* has impaired proliferation and survival within J774 murine macrophages *in vitro* (21). The *Δplb1* strain has a deletion in the *PLB1* gene which codes for the secreted enzyme phospholipase B1 (23). The *Δplb1* strain is known to produce lower levels of fungal eicosanoids indicating that phospholipase B1 is involved in fungal eicosanoid synthesis (19). It has been proposed that the attenuation of this strain within macrophages could be because it cannot produce eicosanoids (19). A previous study has identified PGE_2_ as an eicosanoid that promotes cryptococcal virulence and manipulates macrophage activation, however this study did not determine if PGE_2_ was produced by the host or *C. neoformans* (20). We hypothesised that PGE_2_ - or other phospholipase B1 derived eicosanoid species - are produced by *C. neoformans* during infection and promote macrophage infection.

To test if PGE_2_ promotes the intracellular growth of *C. neoformans* we treated *Δplb1* infected J774 macrophages with exogenous PGE_2_ and measured intracellular proliferation over 18 hours. The addition of exogenous PGE_2_ to J774 macrophages infected with *Δplb1* was sufficient to recover the intracellular proliferation of *Δplb1* compared to the H99 (parental wild type strain) and *Δplb1:PLB1* (reconstituted strain) strains (Fig 1 p= 0.038). These findings support our initial hypothesis and identify PGE_2_ as a mediator of cryptococcal virulence during macrophage infection.

**Fig 1.**
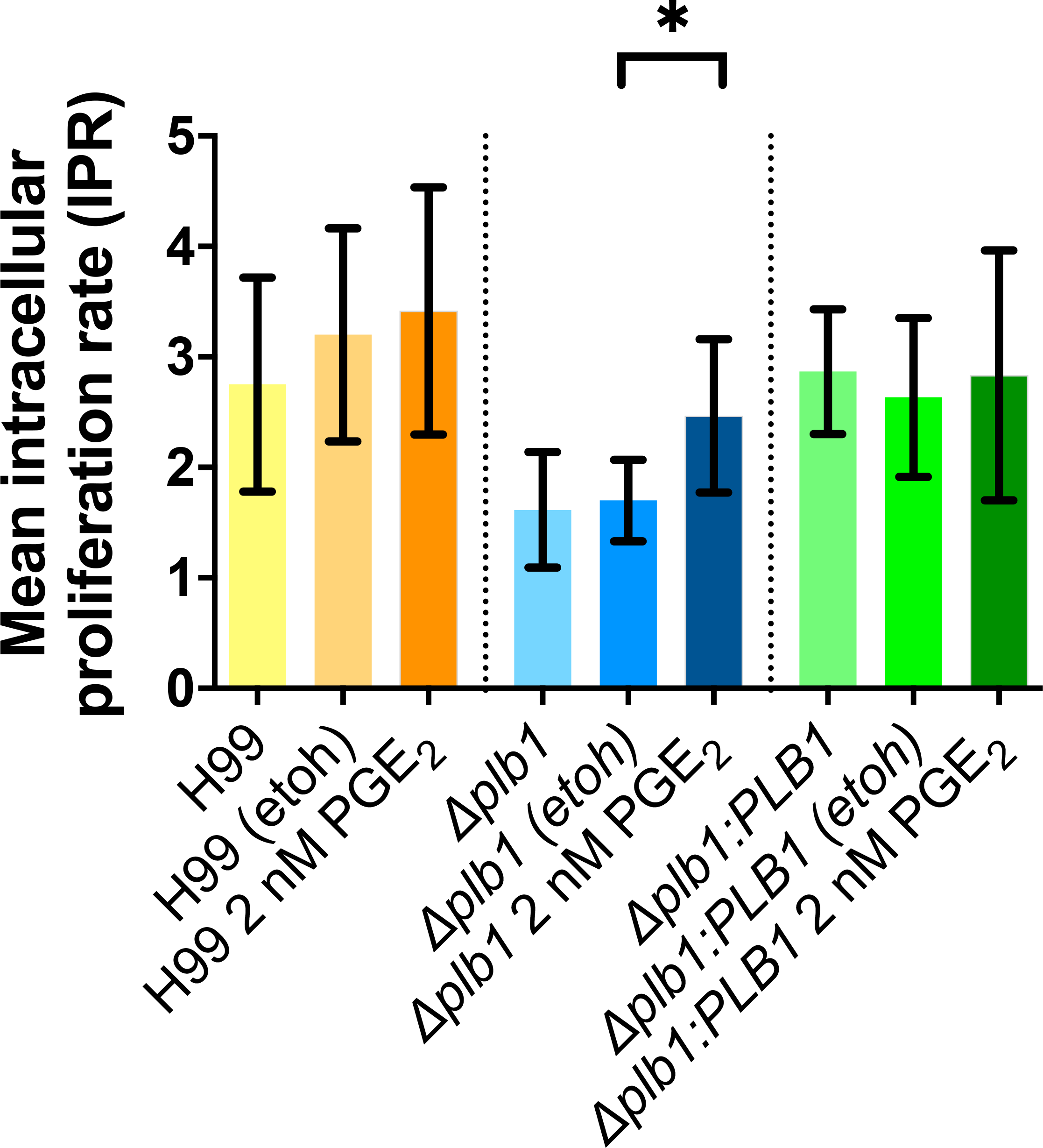
The intracellular proliferation defect of the *C.* neoformans mutant *Δplb1*, can be reversed with the addition of exogenous prostaglandin E_2_. J774 murine macrophages were infected with *plb1,* the parental strain H99 or a genetic reconstitute strain *Δplb1:PLB1*. Infected cells were left untreated or treated with 2 nM PGE_2_ or an equivalent solvent (ethanol) control. Mean IPR from 5 biological repeats shown with error bars representing standard deviation. An unpaired two tailed Student’s t-test was performed to compare each treatment group. H99 etoh vs H99 2 nM PGE_2_ ns p = 0.7212, *Δplb1* etoh vs. *Δplb1* 2 nM PGE_2_ * p = 0.0376, *plb1:PLB1* etoh vs. *Δplb1:PLB1* 2 nM PGE_2_ ns p=0.723.

We have previously shown that *Δplb1* has reduced intracellular proliferation due to a combination of reduced cell division and reduced viability within the phagosome (21). In our intracellular proliferation assay the number of intracellular *Cryptococcus* cells is quantified by lysing infected macrophage and counting the number of *Cryptococcus* cells with a hemocytometer. Due to their structurally stable fungal cell wall dead or dying *Cryptococcus* cells do not look noticeably different on a hemocytometer from viable cells. To quantifiy the viability of *Cryptococcus* cells retrieved from the phagosome we diluted the lysate to give an expected number of CFUs (in this case 200 CFU), spread the diluted lysate on YPD agar and count the actual number of CFUs produced – a difference between the expected CFU count (200 CFU) and the actual CFU count indicates a loss of *Cryptococcus* cell viability. In this case viability assays showed that exogenous PGE_2_ produced no significant increase in the viability of *Δplb1* cells within the phagosome (Supplementary Fig 1A).

### Exogenous prostaglandin E_2_ rescues in vivo growth of Δplb1-GFP

Our *in vitro* data showed that PGE_2_ promotes the intracellular proliferation of *C. neoformans* within macrophages. First, we injected 2-day post fertilisation (dpf) zebrafish larvae with *Δplb1-*GFP (a constitutively expressed GFP tagged version of the *Δplb1* generated for this study). One of the advantages of this model is that the fungal burden can be non-invasively imaged within infected larvae using fluorescently tagged *C. neoformans* strains and we able measure the growth of the fungus at 1-, 2- and 3-days post-infection (dpi). We found that *Δplb1-*GFP infected larvae had significantly lower fungal burdens at 1, 2- and 3-days post infection (Fig 2D and Supplementary Fig 1B) compared to the parental strain H99-GFP (Fig 2A and Supplementary Fig 1B). These data demonstrated that the *Δplb1-*GFP mutant had a similar growth deficiency to our *in vitro* phenotype and with previous studies (21,23,24). To confirm that PGE_2_ promotes cryptococcal infection *in vivo* we infected zebrafish larvae with *Δplb1-*GFP or H99-GFP and treated the larvae with exogenous PGE_2_. In agreement with our *in vitro* findings (Fig 1), exogenous PGE_2_ increased the growth of both the parental H99 strain (Fig 2B, p = 0.0137, 1.35-fold increase vs. DMSO) and the *Δplb1-*GFP mutant (Fig 2E, p= 0.0001, 2.15-fold increase vs. DMSO) while PGD_2_ did not (Fig 2C and 2F). Taken together these data show that PGE_2_ is sufficient to enhance the virulence of *C. neoformans in vivo*, furthermore our *in vitro* data suggest that this is a result of uncontrolled intracellular proliferation within macrophages (Fig 1).

**Fig 2.**
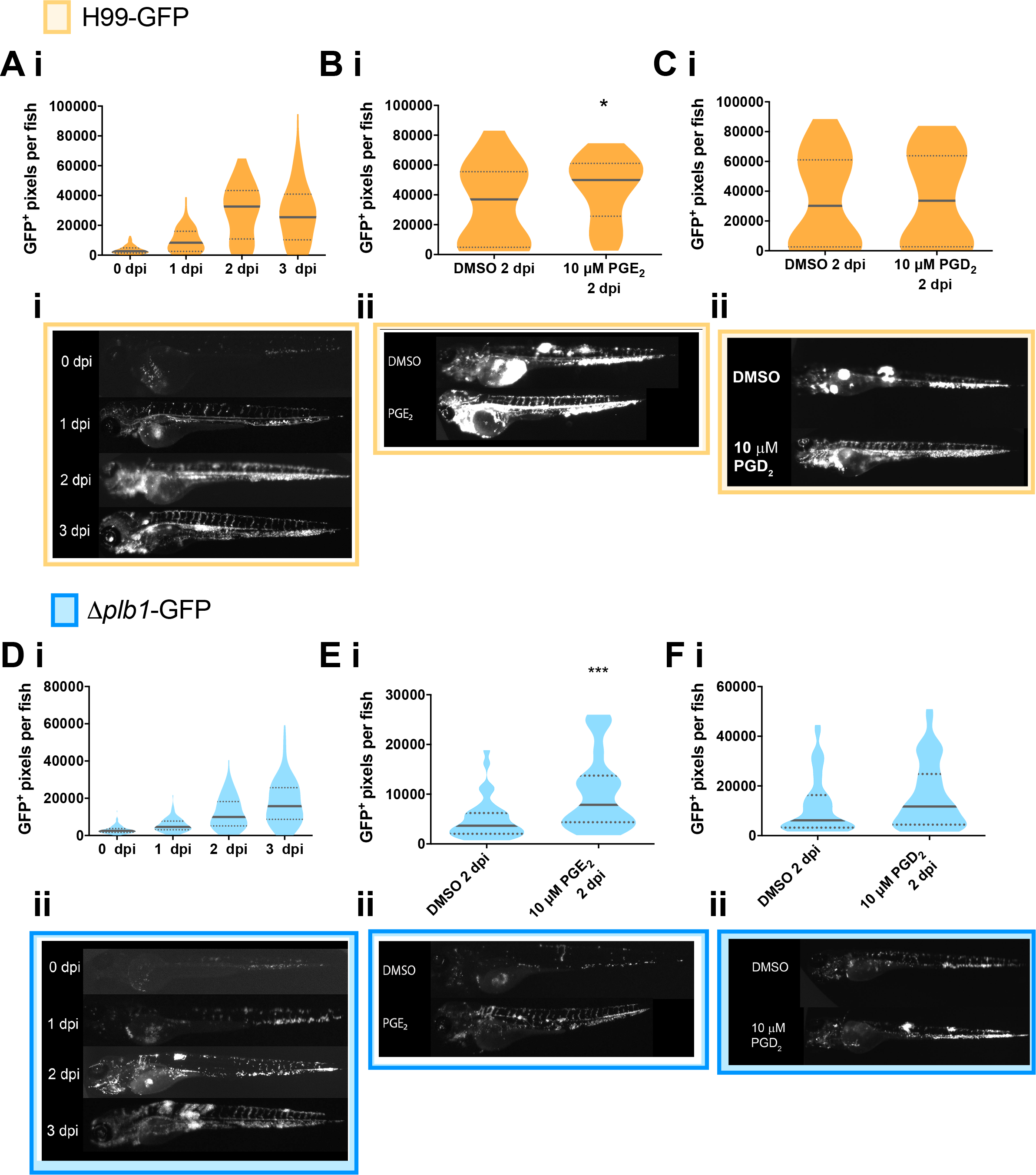
The prostaglandin E_2_ dependent growth defect of *Δplb1* is also present *in vivo*. **A i** H99-GFP infected larvae imaged at 0, 1, 2 and 3 dpi. At least 50 larvae measured per time point across 3 biological repeats. Box and whiskers show median, 5^th^ percentile and 95^th^ percentile. Unpaired Mann-Whitney U tests used to compare the burden between each strain for every time point, for p values see (Supplementary Fig 1B ii). **B i**– H99-GFP Infected larvae treated with 10 μM prostaglandin E_2_ or equivalent solvent (DMSO) control. At least 60 larvae measured per treatment group from 3 biological repeats. Box and whiskers show median, 5^th^ percentile and 95^th^ percentile. Unpaired Mann-Whitney U tests used to compare between treatments DMSO vs. 10 μM PGE_2_ * p = 0.0137 (threshold for significance 0.017, corrected for multiple comparisons). **C i**, H99-GFP Infected larvae treated with 10 μM prostaglandin D_2_ or equivalent solvent (DMSO) control. At least 60 larvae measured per treatment group from 4 biological repeats. Box and whiskers show median, 5^th^ percentile and 95^th^ percentile. Unpaired Mann-Whitney U tests used to compare between treatments DMSO vs. 10 μM PGD_2.,_ ns p= 0.8 **D i***Δplb1-*GFP infected larvae (500 cell inoculum injected at 2 dpf) imaged at 0, 1, 2 and 3 dpi. N = 3. Box and whiskers show median, 5^th^ percentile and 95^th^ percentile. At least 87 larvae measured for each time point from 3 biological repeats. Unpaired Mann-Whitney U tests used to compare the burden between each strain for every time point, for p values see (Supplementary Fig 1 A i + A ii). **E i** *Δplb1-*GFP Infected larvae treated with 10 μM prostaglandin E_2_ or equivalent solvent (DMSO) control. At least 35 larvae measured per treatment group from 2 biological repeats. Box and whiskers show median, 5^th^ percentile and 95^th^ percentile. Unpaired Mann-Whitney U tests used to compare between treatments *Δplb1-*GFP DMSO vs 10 μM PGE_2_ *** p = 0.0001 (threshold for significance 0.017, corrected for multiple comparisons). **F i** *Δplb1*-GFP Infected larvae treated with 10 μM prostaglandin D_2_ or equivalent solvent (DMSO) control. At least 45 larvae measured per treatment group from 3 biological repeats. Box and whiskers show median, 5^th^ percentile and 95^th^ percentile. Unpaired Mann-Whitney U tests used to compare between treatments DMSO vs. 10 μM PGD_2._ Ns p = 0.1 **A ii** Representative GFP images (representative = median value) H99-GFP infected larvae, untreated at 0,1,2,3 dpi. **B ii, C ii** Representative GFP images (representative = median value) H99-GFP infected larvae, at 2 dpi treated with 10 μM PGE_2_ (**B ii**) or PGD_2_ (**C ii**). **D ii** Representative GFP images (representative = median value) *Δplb1-*GFP infected larvae, untreated at 0,1,2,3 dpi. **E ii, F ii** Representative GFP images (representative = median value) *Δplb1-*GFP infected larvae, at 2 dpi treated with 10 μM PGE_2_ (**E ii**) or PGD_2_ (**F ii**).

### Prostaglandin E_2_ must be dehydrogenated into 15-keto-PGE_2_ to promote C. neoformans growth

PGE_2_ can be enzymatically and non-enzymatically modified in cells to form a number of distinct metabolites. To distinguish the biological activity of PGE_2_ rather than its metabolites we used an analogue of PGE_2_ called 16,16-dimethyl PGE_2_ that cannot be dehydrogenated but otherwise has comparable activity to PGE_2_ (25)). We found that unlike PGE_2_, 16,16-dimethyl PGE_2_ treatment did not increase the fungal burden of *Δplb1-*GFP (Fig 3C p= 0.9782) *or* H99-GFP infected larvae (Fig 3A p= 0.9954). Therefore, the biological activity of PGE_2_ alone did not appear to promote cryptococcal pathogenesis and that dehydrogenation of PGE_2_ was required. PGE_2_ and 16,16-dimethyl PGE_2_ both signal through PGE_2_ receptors (EP1, EP2, EP3 and EP4) but 15-keto-PGE_2_ does not. In a murine model of pulmonary cryptococcosis the PGE_2_ receptors EP2 and EP4 have been identified as promoters of fungal virulence (20). To confirm that PGE_2_ itself does not promote virulence in our model we treated zebrafish with antagonists against the EP2 and EP4 receptors. In support of our previous experiment we found that EP2 / EP4 inhibition had no effect on fungal burden in zebrafish (Fig 3E). An abundant dehydrogenated form of PGE_2_ is 15-keto-PGE_2_ (that has also been isolated from *C. neoformans* (18)) and we tested if 15-keto-PGE_2_ was sufficient to rescue the growth defect of the *Δplb1* mutant during infection. Therefore, we treated infected zebrafish larvae with exogenous 15-keto-PGE_2_ and found that this was sufficient to significantly increase the fungal burden of zebrafish larvae infected with both *Δplb1-*GFP (Fig 3D, p = 0.0119, 1.56-fold increase vs. DMSO) and H99-GFP (Fig 3B, p = 0.0048, 1.36-fold increase vs. DMSO). To explore the *Cryptococcus* eicosanoid synthesis pathway further we used a second eicosanoid deficient *C. neoformans* mutant *Δlac1*. Whereas *Δplb1* is unable to produce any eicosanoid species *Δlac1* is deficient only in PGE_2_ and 15-keto-PGE_2_ (18). Using a *Δlac1-*GFP strain generated for this study we found that *Δlac1-*GFP also produced low fungal burden during *in vivo* zebrafish larvae infection (Supplementary Fig 2B). As with *Δplb1-*GFP, this defect could be rescued with the addition of exogenous PGE_2_ (Supplementary Fig 2C), but not with 16,16-dm-PGE_2_ (Supplementary Fig 2D) or with 15-keto-PGE_2_ (Supplementary Fig 2E). Prostaglandin E_2_ is known to affect haematopoietic stem cell homeostasis in zebrafish (26). This could affect macrophage number and subsequently fungal burden. We have previously observed that a large depletion of macrophages can lead to increased fungal burden in zebrafish larvae (22). We performed whole body macrophage counts on 2 dpf uninfected larvae treated with PGE_2_ or 15-keto-PGE_2_ 2 days post treatment (the same time points used in our infection assay). Following PGE_2_ and 15-keto PGE_2_ treatment macrophages were still observable throughout the larvae. For PGE_2_ treatment we saw on average a 15% reduction in macrophage number while 15-keto PGE_2_ did not cause any decrease (Supplementary Fig 1C). Due to the fact that fungal burden increased during both PGE_2_ and 15-keto PGE_2_ treatments it is highly unlikely that this reduction could account for the increases in burden seen.

**Fig 3.**
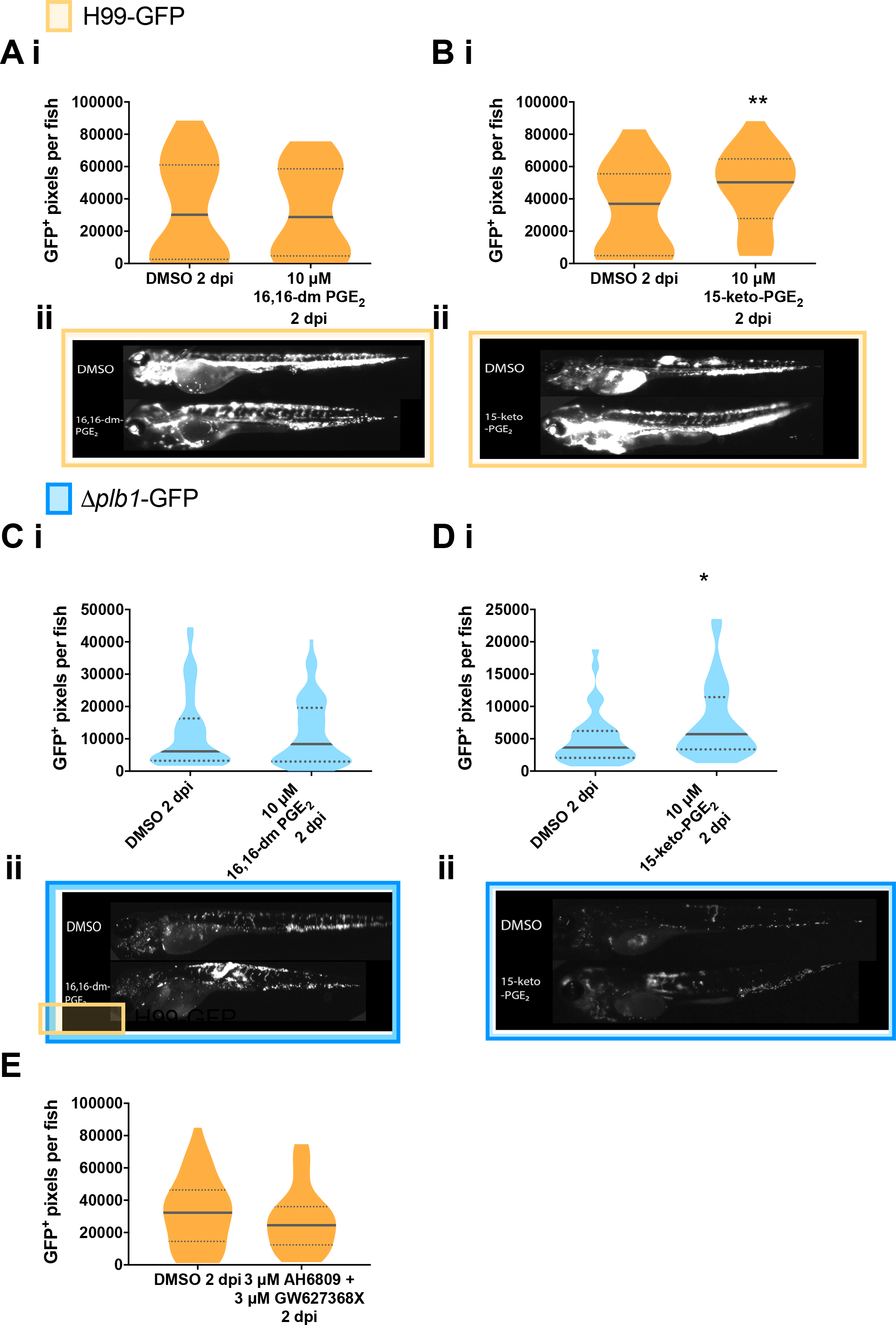
The observed activity of PGE_2_ is due to its dehydrogenated derivative 15-keto-PGE_2_. Fungal burden measured at 2 days post infection (2 dpi) by counting GFP positive pixels in each larvae **A i** H99-GFP Infected larvae treated with 10 μM 16,16-dimethyl-prostaglandin E_2_ or equivalent solvent (DMSO) control. At least 75 larvae measured per treatment group from 4 biological repeats. Box and whiskers show median, 5^th^ percentile and 95^th^ percentile. Unpaired Mann-Whitney U test used to compare between treatments, DMSO vs. 10 μM 16, 16-dm PGE_2_ ns p = 0.9954. **B i** H99-GFP Infected larvae treated with 10 μM 15-keto-prostaglandin E_2_ or equivalent solvent (DMSO) control. At least 55 larvae measured per treatment group from 3 biological repeats. Unpaired Mann-Whitney U test used to compare between treatments DMSO vs. 10 μM 15-keto=PGE_2_ ** p = 0.0048 (threshold for significance 0.017, corrected for multiple comparisons). **C i** *Δplb1-*GFP Infected larvae treated with 10 μM 16, 16-dimethyl prostaglandin E_2_ or equivalent solvent (DMSO) control. At least 45 larvae per treatment group from 3 biological repeats. Unpaired Mann-Whitney U test used to compare between treatments *Δplb1-*GFP DMSO vs 10 μM 16, 16-dm PGE_2_ ns p = 0.98. **D i** *Δplb1-*GFP Infected larvae treated with 10 μM 15-keto-prostaglandin E_2_ or equivalent solvent (DMSO) control. At least 35 larvae measured per treatment group from 2 biological repeats. Unpaired Mann-Whitney U test used to compare between treatments DMSO vs 10 μM 15-keto-PGE_2_ * p = 0.0119 (threshold for significance 0.017, corrected for multiple comparisons). **A ii, B ii** Representative GFP images (representative = median value) H99-GFP infected larvae, at 2dpi treated with 10 μM 16,16-dm-PGE_2_ (**A ii**) or 15-keto-PGE_2_ (**B ii**). **C ii, D ii** Representative GFP images (representative = median value) *Δplb1-*GFP infected larvae, at 2dpi treated with 10 μM 16,16-dm-PGE_2_ (**C ii**) or 15-keto-PGE_2_ (**D ii**). **E** H99-GFP infected larvae treated with a combination of 3 μM AH6809 and 3 μM GW627368X or equivalent solvent (DMSO) control. Box and whiskers show median, 5^th^ percentile and 95^th^ percentile. At least 64 larvae measured per treatment group from 4 biological repeats. Mann-Whitney U test used to compare between treatments, no significance found.

### Host derived prostaglandins are not required for growth of C. neoformans

After determining that PGE_2_ promotes the growth of *C. neoformans in vitro* and *in vivo* via its metabolite 15-keto-PGE_2_, we wanted to determine whether the source of these prostaglandins was the host or the fungus. Our *in vitro* data show that the *C. neoformans* strain *Δplb1* has a growth deficiency *in vitro* and *in vivo* that can be rescued with the addition of PGE_2_, because this phenotype is mediated by cryptococcal phospholipase B1 it indicates that pathogen-derived rather than host-derived prostaglandins are required. A previous study reports that *C. neoformans* infection can induce higher PGE_2_ levels in the lung during *in vivo* pulmonary infection of mice (20), although it was not determined if the PGE_2_ was host or pathogen derived. Therefore, we tested the hypothesis that host prostaglandin synthesis was not required for cryptococcal virulence.

To block host prostaglandin synthesis *in* vitro we inhibited host cyclooxygenase activity because it is essential for prostaglandin synthesis in vertebrates (27). We treated H99 and *Δplb1* infected J774 macrophages with aspirin at a concentration we determined was sufficient to block host PGE_2_ synthesis (Supplementary Fig 3A). Aspirin is a non-reversible inhibitor of cyclooxygenase 1 (COX-1) and cyclooxygenase 2 (COX-2) enzymes, we therefore included a condition where J774 cells were pretreated with aspirin only prior to infection and then at a condition where aspirin was present throughout infection. We found that aspirin treatment did not affect the intracellular proliferation of H99 or *Δplb1* (Fig 4A), suggesting that host cyclooxygenase activity is not required for the phospholipase B1 dependent virulence of *C. neoformans* during macrophage infection. To confirm these *in vivo,* we pharmacologically blocked zebrafish cyclooxygenase-1 (COX-1) and cyclooxygenase-2 (COX-2). We used separate cyclooxygenase inhibitors with zebrafish larvae instead of aspirin because we found that aspirin treatment led to lethal developmental defects in zebrafish larvae (unpublished observation). We infected 2 dpf zebrafish larvae with H99-GFP and *Δplb1-*GFP and treated with inhibitors for COX-1 (NS-398, 15 μM) and COX-2 (SC-560, 15 μM). We found that both inhibitors decreased the fungal burden of H99-GFP, but not *Δplb1-*GFP infected zebrafish larvae (Fig 4B i and 4B ii, H99-GFP - NS-398, p= 0.0002, 1.85-fold decrease vs. DMSO. SC-560 p= <0.0001, 3.14-fold decrease vs DMSO). These findings were different to what we had observed *in vitro* but because this phenotype was phospholipase B1 dependent we reasoned that these inhibitors could be having off target effects on *C. neoformans* - although *C. neoformans* does not have a homolog to cyclooxygenase other studies have tried to inhibit eicosanoid production in *Cryptococcus* using cyclooxygenase inhibitors but their efficacy and target remain uncertain (17,28). To support our pharmacological evidence, we used a CRISPR/Cas9-mediated knockdown of the prostaglandin E_2_ synthase gene (*ptges)* (29). We used a knockdown of tyrosinase (*tyr*) – a gene involved in the conversion of tyrosine into melanin as a control because *tyr*^−/−^crispants are easy to identify because they do not produce any pigment. We infected 2 dpf *ptges*^−/−^and *tyr*^−/−^zebrafish larvae with H99-GFP or *Δplb1-*GFP and measured the fungal burden at 3 dpi. We found that *ptges*^−/−^ zebrafish infected with H99-GFP had a higher fungal burden at 3 dpi compared to *tyr*^−/−^zebrafish infected with H99-GFP whereas there was no difference between *ptges*^−/−^ and *tyr*^−/−^zebrafish larvae infected with *Δplb1-*GFP (Fig 4C). Thus, both pharmacological and genetic inhibitions of host prostaglandin synthesis were not determinants of C. neoformans growth.

**Fig 4.**
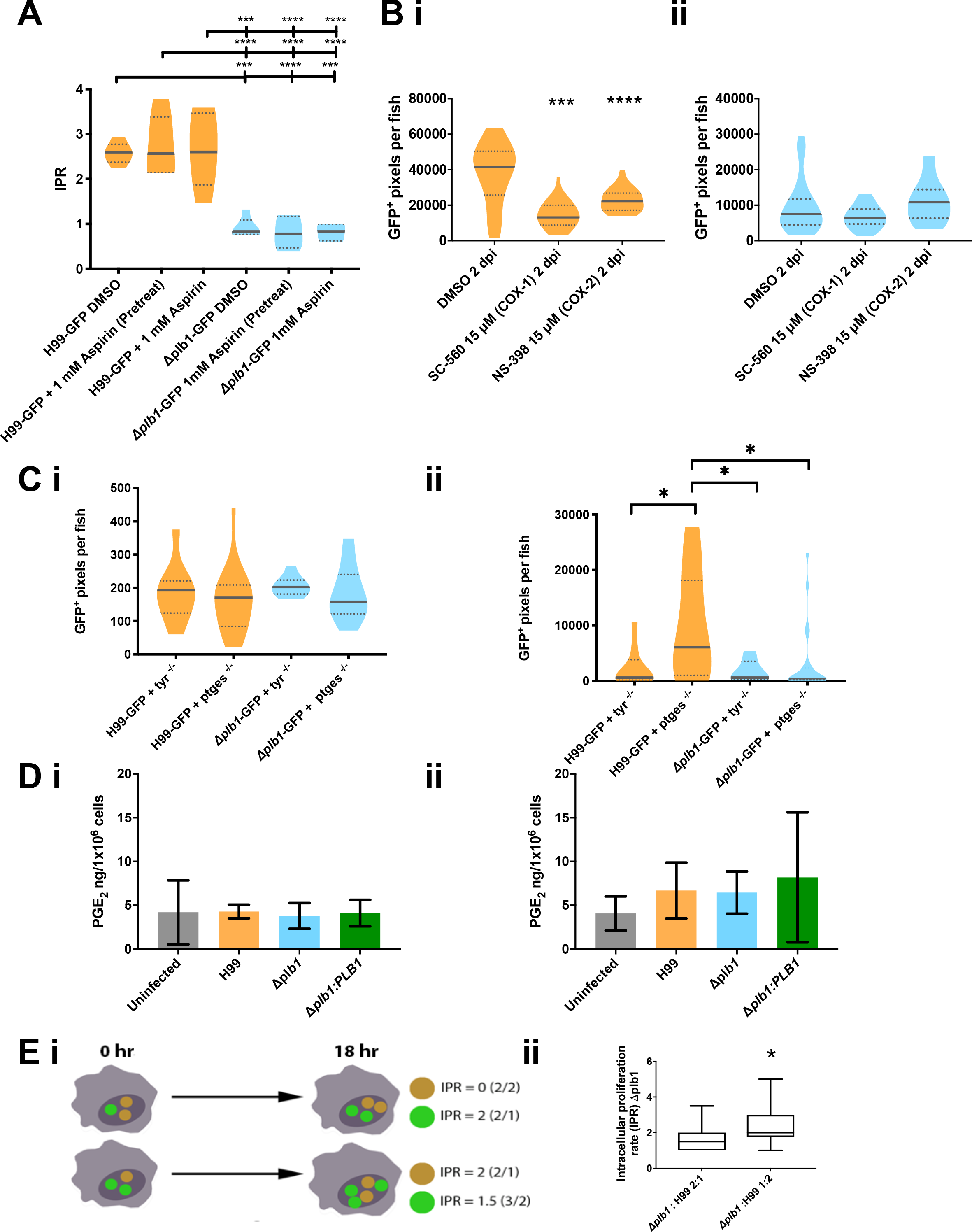
Host derived prostaglandins are not required for growth of C. neoformans. A Intracellular proliferation quantified from timelaspe movies of J774 macrophages infected with H99-GFP or *Δplb1-*GFP and treated with 1 mM Aspirin – either for 18 hours before infection (pretreatment) or throughout the time course of infection. One-way ANOVA with Tukey post-test performed comparing all conditions. H99-GFP DMSO vs. *Δplb1-*GFP DMSO *** p = 0.0002. H99-GFP DMSO vs. *Δplb1-*GFP 1 mM (pretreat) **** p = <0.0001. H99-GFP DMSO vs. *Δplb1-*GFP 1 mM aspirin *** p = 0.0001. H99-GFP + 1mM aspirin (pretreat) vs. *Δplb1-*GFP DMSO **** p <0.0001. H99-GFP + 1mM aspirin (pretreat) vs. *Δplb1-*GFP 1 mM aspirin (pretreat) **** p < 0.0001. H99-GFP + 1mM aspirin (pretreat) vs. *Δplb1-*GFP 1 mM aspirin **** p <0.0001. H99-GFP+ 1mM aspirin vs. *Δplb1-*GFP DMSO *** p = 0.0001. H99-GFP + 1mM aspirin vs. *Δplb1-*GFP 1 mM aspirin (pretreat) **** p <0.0001. H99-GFP+ 1mM aspirin vs. *Δplb1-*GFP 1 mM aspirin **** p < 0.0001. **B i** H99-GFP infected larvae treated with 15 μM NS-398, 15 μM SC-560 or equivalent solvent (DMSO) control. Box and whiskers show median, 5^th^ percentile and 95^th^ percentile. At least 25 larvae measured per treatment group from 2 biological repeats. Mann-Whitney U test used to compare between treatments, DMSO vs. 15 μM *** p = 0.0002. **B ii** *Δplb1-*GFP infected larvae treated with 15 μM NS-398, 15 μM SC-560 or equivalent solvent (DMSO) control. Box and whiskers show median, 5^th^ percentile and 95^th^ percentile. At least 34 larvae measured per treatment group from 2 biological repeats. Mann-Whitney U test used to compare between treatments, no significance found. **C**2 dpi zebrafish larvae crispants with CRISPR knockout against Prostaglandin E_2_ synthase (*ptges)* or a Tyrosinase control (*tyr*) infected with H99-GFP or *Δplb1-*GFP, fungal burden quantified at 0 dpi (**C i**) and 3 dpi (**C ii**) – data shown is a single experiment but is representative of N=3 experiments. One way ANOVA with Tukey post-test performed to compare each condition **C i** no significance found **C ii** H99-GFP + *tyr* ^−/−^ vs. H99-GFP + *ptges* ^−/−^ * p = 0.0390. H99-GFP + *ptges* ^−/−^ vs. *Δplb1-*GFP + *tyr* ^−/−^ * p = 0.0313. H99-GFP + *ptges* ^−/−^ vs. *Δplb1-*GFP + *ptges* ^−/−^ * p = 0.0121. **D i** PGE_2_ monoclonal EIA ELISA performed on supernatants from *C. neoformans* infected macrophages collected at 18 hr post infection. Mean concentration of PGE_2_ (pg per 1×10^6^ cells) plotted with SD, n = 4. One-way ANOVA with Tukey post-test performed, no significance found. **D ii** LC MS/MS mass spectrometry analysis performed on cell suspensions (infected J774 cells and supernatants) collected at 18 hr post infection. Mean concentration of PGE_2_ (pg per 1×10^6^ cells) plotted with SD, n = 3. One-way ANOVA with Tukey post-test performed, no significance found. **E** J774 cells co-infected with a 50:50 mix of *Δplb1* and H99-GFP. **i** Diagrammatic representation of co-infection experiment. GFP^+^ (green) and GFP^−^ (yellow) *C. neoformans* cells within the phagosome were quantified at 0 hr, macrophages with a burden ratio of 1:2 or 2:1 were re-analysed at 18 hr, the IPR for *Δplb1* within 2:1 and 1:2 co-infected cells were calculated by dividing the burden at 18hr by burden at 0 hr for GFP^+^ (green) or GFP^−^ (yellow) cells. **ii** Quantification of IPR for *Δplb1* cells within *Δplb1*:H99-GFP 2:1 or 1:2 co-infected macrophages. At least 35 co-infected macrophages were analysed for each condition over 4 experimental repeats. Student’s T test performed to compare ratios – 2:1 vs 1:2 * p = 0.0137.

### Phospholipase B1 dependent factors are sufficient to support Δplb1 growth in macrophages

To further evidence that *C. neoformans* was the source of PGE_2_ during macrophage infection we used a co-infection assay which has previously been used to investigate the interaction of different *C. gattii* strains within the same macrophage (30). We hypothesised that if *C. neoformans* derived prostaglandins promoted fungal growth, co-infection between H99 and *Δplb1* would support mutant growth within macrophages because the parental strain H99 would produce growth promoting prostaglandins that are lacking in *Δplb1*. To produce co-infection, J774 murine macrophages were infected with a 50:50 mixture of *Δplb1* and H99-GFP (31) (Fig 4E i; as described previously for *C. gattii* (30)). This approach allowed us to differentiate between *Δplb1* (GFP negative) and H99 (GFP positive) *Cryptococcus* strains within the same macrophage and to score their proliferation separately. The intracellular proliferation of *Δplb1* was calculated by counting the change in number of GFP negative *Δplb1* cells over an 18hr period from time-lapse movies of infected cells. We found that co-infected macrophages did not always contain an equal ratio of each strain at the start of the 18hr period so we scored the proliferation of *Δplb1* for a range of initial burdens (1:2, 1:3 and 1:4). We found that *Δplb1* proliferated better when accompanied by two H99-GFP yeast cells in the same macrophage (Fig 4E ii, 1:2 p = 0.014) as opposed to when two *Δplb1* yeast cells were accompanied by one H99-GFP yeast cell (Fig 4E ii, 2:1). We observed a similar effect for ratios of 1:3 and 1:4, but these starting burden ratios are particularly rare, under powering our analysis (Supplementary Fig 3). This effect was also recapitulated for J774 macrophages co-infected with *Δlac1* and H99-GFP (Supplementary Fig 2A). These data indicate that a phospholipase B1 dependent factor found in H99 infected macrophages, but absent in *Δplb1* and *Δlac1* infected macrophages, is required for intracellular proliferation within macrophages.

### Production of PGE_2_ by macrophages is not altered by C. neoformans infection

Our data indicate that host cells are not the source of virulence promoting prostaglandins, and that secreted factors produced by wild type - but not *Δplb1* or *Δlac1 Cryptococcus* strains – promote fungal growth within macrophages. We wanted to test if there were detectable difference in PGE_2_ levels between infected and uninfected macrophages caused by cryptococcal prostaglandin synthesis. To do this we performed ELISA analysis to detect PGE_2_ concentrations in supernatants from *C. neoformans* infected J774 macrophages (Fig 4D i). We found that J774 macrophages produced detectable levels of PGE_2_ (mean concentration 4.10 ng/1×10^6^ cells), however we did not see any significant difference between infected or uninfected macrophages (infected with H99, *Δplb1* or *Δplb1:PLB1* strains). To confirm our ELISA results, we performed LC MS/MS analysis of lysed J774 macrophages using a PGE_2_ standard for accurate quantification (Fig 4D ii). The concentrations detected were similar to those measured by our ELISA (mean concentration 6.35 ng/1×10^6^ cells) and also did not show any significant differences between conditions. Taken together these data suggest that any *Cryptococcus*-derived prostaglandins present during infection were likely to be contained within the macrophage in low, localized concentrations and that the host receptor targeted by these eicosanoids is therefore likely to be intracellular.

### 15-keto-PGE_2_ promotes C. neoformans growth by activating host PPAR-γ

We next wanted to determine how PGE_2_ / 15-keto-PGE_2_ promotes *C. neoformans* infection. We hypothesized that these prostaglandins were interfering with inhibition of fungal growth via a host receptor. Our experiments inhibiting EP2 / EP4 suggest that 15-keto-PGE_2_ must signal through a different receptor to PGE_2_ (Fig 3E). 15-keto-PGE_2_ is a known agonist of the peroxisome proliferation associated receptor gamma (PPAR-*γ*) (32); a transcription factor that controls expression of many inflammation related genes (33–35). We first tested if PPAR-*γ* activation occurs within macrophages during *C. neoformans* infection by performing immunofluorescent staining for PPAR-*γ* in macrophages infected with H99 and *Δplb1*. To quantify PPAR-*γ* activation we measured its nuclear translocation by comparing nuclear and cytoplasmic fluorescence intensity in infected cells. PPAR-*γ* is a cytosolic receptor that translocates to the nucleus upon activation, therefore cells where PPAR-*γ* is activated should have increased nuclear staining for PPAR-*γ*. We found that J774 macrophages infected with H99 had significantly higher levels of nuclear staining for PPAR-*γ* compared to *Δplb1* infected and uninfected cells (Fig 5A i and Supplementary Fig 3C). This confirmed that *C. neoformans* activates PPAR-*γ* and that this phenotype is phospholipase B1 dependent.

**Fig 5.**
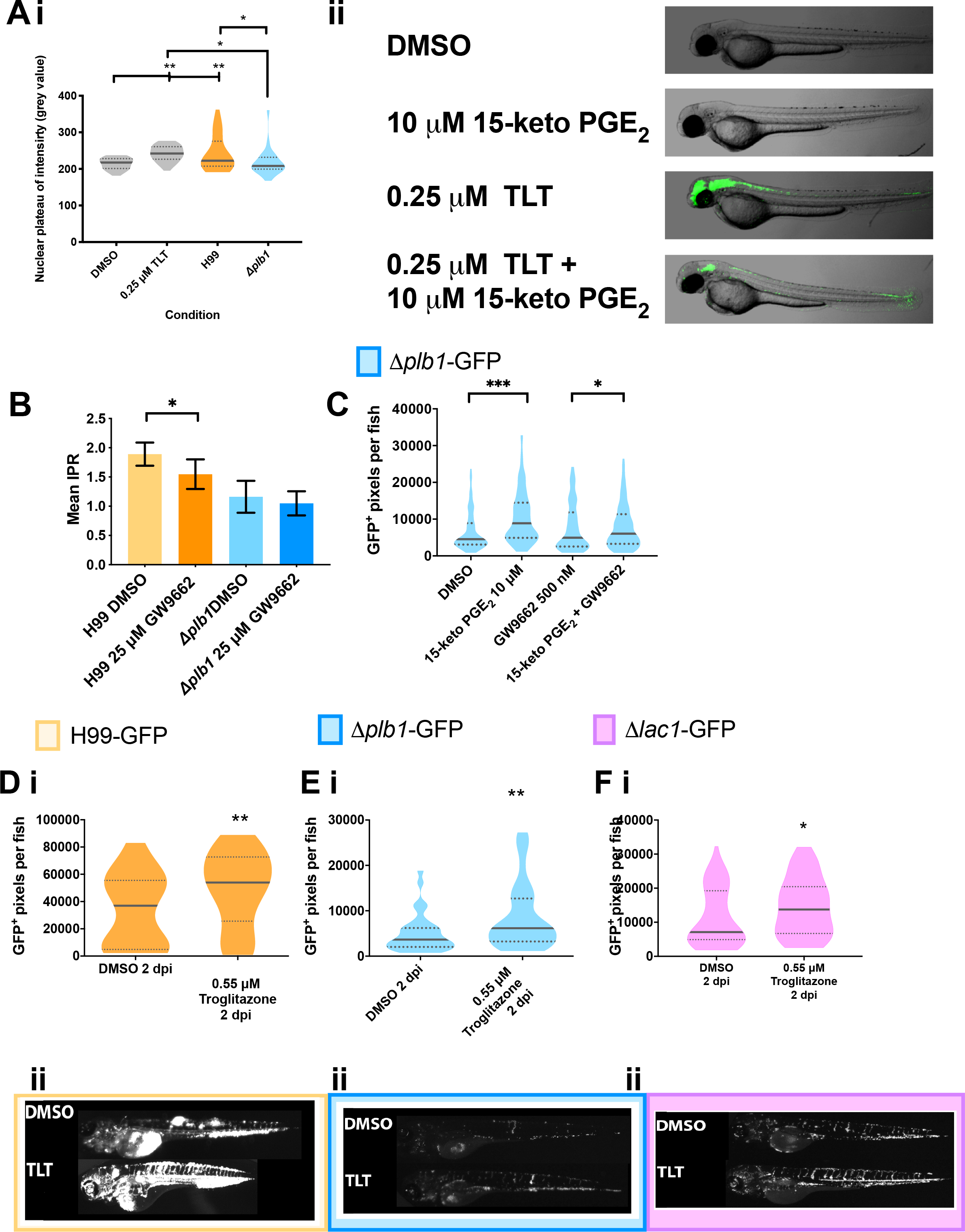
15-keto-PGE_2_ promotes fungal burden by activating host PPAR-*γ*. **A i** J774 macrophages treated with Troglitazone (TLT - 0.25 μM), an equivalent DMSO control or infected with H99 or *Δplb1* fixed and stained at 18 hpi with Hoechst and antibody against PPAR-γ Nuclear localization of PPAR-γ quantified by measuring nuclear grey value of at least 30 cells per condition. A single experiment is shown that is representative of n = 2. One way ANOVA with Tukey post-test used to compare all conditions. DMSO vs. TLT ** p = 0.0049. DMSO vs H99 ** p = 0.0040. TLT vs *Δplb1* * p = 0.0212. H99 vs. *Δplb1* * p = 0.0187. **A ii** Transgenic zebrafish larvae with a GFP PPAR- γ reporter treated with DMSO, 250 nM Troglitazone or 10 μM 15-keto-PGE_2_ + 250 nM Troglitazone overnight and imaged. Lateral views of 2 dpf embryos, anterior to the left, are shown. **B** J774 murine macrophages infected with *Δplb1* or the parental strain H99. Infected cells treated with 25 μM GW9662 (a PPAR-γ antagonist) or equivalent solvent (DMSO) control. Mean IPR from 6 biological repeats shown with error bars representing standard deviation. An unpaired two tailed Student’s t-test was performed to compare each treatment group. H99 DMSO vs. H99 25 μM GW9662 * p = 0.026. **C** Δplb1-GFP infected larvae treated with 10 μM 15-keto-PGE_2_, 500 nM GW9662, 10 μM 15- keto-PGE_2_ + 500 nM GW9662 or an equivalent solvent (DMSO) control. Box and whiskers show median, 5^th^ percentile and 95^th^ percentile. At least 35 larvae measured per treatment group from 2 biological repeats. Mann-Whitney U test used to compare between treatments. DMSO vs. 15- keto-PGE_2_ **** p <0.0001 (threshold for significance 0.025, corrected for multiple comparisons), 15-keto-PGE_2_ vs. 15-keto-PGE_2_ + 500 nm GW9662 ** p = 0.005 (threshold for significance 0.025, corrected for multiple comparisons) **D-F** 2 day old (2 dpf) *Nacre* zebrafish larvae injected with 500 cell inoculum. Fungal burden measured at 2 days post infection (2 dpi) by counting GFP positive pixels within each larvae. **D i** H99-GFP Infected larvae treated with 0.55 μM Troglitazone (TLT) equivalent solvent (DMSO) control. Box and whiskers show median, 5^th^ percentile and 95^th^ percentile. At least 55 larvae measured per treatment group over 3 biological repeats. Mann-Whitney U test used to compare between treatments, DMSO vs. 0.55 μM Troglitazone ** p = 0.0044 (threshold for significance 0.025, corrected for multiple comparisons). **E i** *Δplb1-*GFP infected larvae treated with 0.55 μM Troglitazone (TLT) equivalent solvent (DMSO) control. Box and whiskers show median, 5^th^ percentile and 95^th^ percentile. At least 35 larvae measured per treatment group from 2 biological repeats. Mann-Whitney U test used to compare between treatments, DMSO vs. 0.55 μM Troglitazone ** p = 0.0089 (threshold for significance 0.025, corrected for multiple comparisons). **F i** *Δlac1-*GFP infected larvae treated with 0.55 μM Troglitazone (TLT) equivalent solvent (DMSO) control. At least 60 larvae measured per treatment group from 3 biological repeats. Box and whiskers show median, 5^th^ percentile and 95^th^ percentile, DMSO vs. 0.55 μM Troglitazone * p = 0.01 (threshold for significance 0.025, corrected for multiple comparisons). **D ii, E ii and F ii**, Representative GFP images (representative = median value) *C. neoformans* infected larvae, at 2 dpi treated with 0.55 μM Troglitazone (TLT) **D ii**-H99-GFP, **E ii** - *Δplb1-*GFP and **F ii** *Δlac1-*GFP.

To test the activation of PPAR-*γ* during infection we first wanted to confirm that exogenous 15-keto-PGE_2_ activates zebrafish PPAR-*γ in vivo* by using transgenic PPAR-*γ* reporter zebrafish larvae (36,37) We treated these larvae at 2 dpf with 15-keto-PGE_2_ and Troglitazone (TLT) which is a specific agonist of PPAR-*γ*. TLT treatment was performed with a concentration (0.55 μM) previously shown to strongly activate PPAR-*γ* in these zebrafish larvae (36). We found that TLT treatment at 2 dpf strongly activated GFP reporter expression in the larvae. We employed a receptor competition assay as a sensitive measurement of binding by simultaneously treating zebrafish with with 15-keto-PGE_2_ and TLT. We observed a reduction in GFP expression compared to TLT treatment alone (Fig 5 Aii), demonstrating competition for the same receptor. This could mean that 15-keto-PGE_2_ was a partial agonist (38,39) or an antagonist to PPAR-*γ* in this experiment. Existing studies suggest 15-keto-PGE_2_ is an agonist to PPAR-*γ* (32) but to confirm this in our model we treated *Δplb1-*GFP infected zebrafish larvae with exogenous 15-keto-PGE_2_ as before but at the same time treated fish with the PPAR-*γ* antagonist GW9662. We found that 15-keto-PGE_2_ treatment significantly improved the growth of *Δplb1-*GFP during infection but that inhibition of PPAR-*γ* was sufficient to reverse this effect (Fig 5C). Therefore, we could demonstrate that that 15-keto-PGE_2_ was an agonist to PPAR-*γ*, and that PPAR-*γ* activation was sufficient to promote a permissive environment for *C. neoformans* growth during infection.

To determine if PPAR-*γ* activation by *C. neoformans* facilitates growth within macrophages we treated H99 and *Δplb1* infected J774 murine macrophages with the PPAR-*γ* antagonist GW9662. GW9662 treatment significantly reduced the proliferation of H99, but not *Δplb1* (Fig 5B, p=0.026, 1.22-fold decrease vs. DMSO). Further supporting that PPAR-*γ* activation is necessary for the successful intracellular parasitism of host macrophages by *C. neoformans*. Finally, to confirm that PPAR-*γ* activation alone promoted *C. neoformans* infection we treated 2dpf infected zebrafish larvae with TLT at the same concentration known to activate PPAR-*γ* in PPAR-*γ* reporter fish (Fig 5A ii and (36)). We found that TLT treatment significantly increased the fungal burden of *Δplb1-*GFP (Fig 5E and, p = 0.0089, 1.68-fold increase vs. DMSO), H99- GFP (Fig 5D, p = 0.0044, 1.46-fold increase vs. DMSO) and *Δlac1-*GFP (Fig 5F, p = 0.01, 1.94-fold increase vs. DMSO) infected larvae similar to 15-keto-PGE_2_ treatment. Thus, we could show that host PPAR-*γ* activation was sufficient to promote cryptococcal growth during infection and was a consequence of fungal derived prostaglandins.

## Discussion

We have shown for the first time that eicosanoids produced by *C. neoformans* promote fungal virulence both *in vitro* and *in vivo*. In this respect we have shown that the intracellular growth defects of two eicosanoid deficient *C. neoformans* strains *Δplb1* and *Δlac1* (21,23) can be rescued with the addition of exogenous PGE_2_. Furthermore, our *in vitro* co-infection assay, *in vitro* infection assays with aspirin, and *in vivo* infection assays provide evidence that the source of this eicosanoid during infection is from the pathogen, rather than the host. Using an *in vivo* zebrafish larvae model of cryptococcosis we find that that PGE_2_ must be dehydrogenated into 15-keto-PGE_2_ before to influence fungal growth. Finally, we provide evidence that the mechanism of PGE_2_/15-keto-PGE_2_ mediated growth promotion during larval infection is via the activation of PPAR-*γ* (33,34,40-42).

In a previous study it was identified that the *C. neoformans* mutant *Δplb1* (that lacks the *PLB1* gene coding for phospholipase B1) was deficient in replication and survival in macrophages (21), a phenotype also observed by a number of studies using different *in vitro* infection assays (19,23). In this study we demonstrate that supplementing *Δplb1* with exogenous prostaglandin E_2_ during *in vitro* macrophages infection is sufficient to restore the mutant’s intracellular proliferation defect. A key goal of this study was to investigate how eicosanoids produced by *C. neoformans* modulate pathogenesis *in vivo* (22). To facilitate *in vivo* measurement of fungal burden we created two GFP-tagged strains with constitutive GFP expression - *Δplb1-*GFP and *Δlac1-*GFP - to use alongside the GFP-tagged H99 parental strain previously produced (31). These two mutants are the only *C. neoformans* mutants known to have a deficiency in eicosanoid synthesis. *Δplb1* cannot produce any eicosanoid species suggesting phospholipase B1 is high in the eicosanoid synthesis pathway while *Δlac1* has a specific defect in PGE_2_ suggesting it might be a prostaglandin E_2_ synthase enzyme. To our knowledge these are the first GFP tagged versions of *Δplb1* and *Δlac1* created. Characterisation of *Δplb1-*GFP and *Δlac1-*GFP *in vivo* revealed that both strains have significantly reduced fungal burdens compared to H99-GFP. This is the first report of *Δlac1* in zebrafish but these observations do confirm a previous zebrafish study showing that non-fluorescent *Δplb1* had attenuated infectious burden in zebrafish larvae (43). To confirm that PGE_2_ is also required for cryptococcal growth *in vivo* we treated *Δplb1-*GFP and *Δlac1-*GFP infected zebrafish larvae with exogenous PGE_2_ to determine how it would affect fungal burden. In agreement with our *in vitro* findings we found that PGE_2_ significantly improved the growth of both of these strains within larvae. Interestingly we also found that PGE_2_ improved the growth of H99-GFP, perhaps representing a wider manipulation of host immunity during *in vivo* infection.

In vertebrate cells, PGE_2_ is converted into 15-keto-PGE_2_ by the enzyme 15-prostaglandin dehydrogenase (15PGDH), furthermore it has been reported that *C. neoformans* has enzymatic activity analogous to 15PGDH (18). To investigate how the dehydrogenation of PGE_2_ to 15- keto-PGE_2_ influenced fungal burden we treated infected larvae with 16,16-dm-PGE_2_ – a synthetic variant of PGE_2_ which is resistant to dehydrogenation (25). Interestingly we found that 16,16-dm-PGE_2_ was unable to promote the growth of *Δplb1-*GFP, H99-GFP or *Δlac1-*GFP within infected larvae. These findings indicate that 15-keto-PGE_2_, rather than PGE_2_, promotes cryptococcal virulence. We subsequently treated infected larvae with exogenous 15-keto-PGE_2_ and confirmed that 15-keto-PGE_2_ treatment was sufficient to promote the growth of both *Δplb1-*GFP and H99-GFP, but not *Δlac1-*GFP (discussed below), without the need for PGE_2_. We therefore propose that PGE_2_ produced by *C. neoformans* during infection must be enzymatically dehydrogenated into 15-keto-PGE_2_ to promote cryptococcal virulence. These findings represent the identification of a new virulence factor (15-keto-PGE_2_) produced by *C. neoformans,* as well as the first-time identification of an eicosanoid other than PGE_2_ with a role in promoting cryptococcal growth. Furthermore, our findings suggest that previous studies which identify PGE_2_ as a promoter of cryptococcal virulence (19,20,44) may have observed additive effects from both PGE_2_ and 15-keto-PGE_2_ activity.

Our experiments with the *Δlac1-*GFP strain reveals that this strain appears to respond in a similar way to PGE_2_, 16,16-dm-PGE_2_ and troglitazone as *Δplb1-*GFP but appears to be unresponsive to 15-keto-PGE_2_. At this time, we cannot fully explain this phenotype, *Δlac1-*GFP was generally less responsive to PGE_2_ and troglitazone treatments compared to *Δplb1-*GFP so it is possible that that higher concentrations of 15-keto-PGE_2_ would be needed to rescue its growth defect, however experimentation with higher concentrations of 15-keto-PGE_2_ led to significant host toxicity. The unresponsiveness of *Δlac1* to eicosanoid treatment could be due to unrelated virulence defects caused by laccase deficiency. Cryptococcal laccase expression is required for the production of fungal melanin – a well characterized virulence factor produced by *C. neoformans* (45,46). It is therefore likely that virulence defects unrelated to eicosanoid synthesis are responsible for the differences between the two mutant phenotypes.

We have found that the phospholipase B1 dependent attenuation of *Δplb1* can be rescued with the addition of exogenous PGE_2_. This indicates that synthesis and secretion of PGE_2_ by *C. neoformans* is a virulence factor. Although our data indicated that *C. neoformans* was the source of PGE_2_ we wanted exclude the possibility that host-derived PGE_2_ was also contributing to virulence. To explore this possibility, we blocked host prostaglandin synthesis - we reasoned that if host PGE_2_ was not required that blocking its production would not affect the growth of *C. neoformans*. To do this we first treated J774 macrophages infected with H99-GFP and *Δplb1-*GFP with aspirin – a cyclooxygenase inhibitor that blocks both COX-1 and COX-2 activity – and found that this had no effect on intracellular proliferation. Subsequently we attempted to block cyclooxygenase activity in zebrafish but found aspirin was lethal at the zebrafish larvae’s currently stage of development, instead we used individual inhibitors specific for COX-1 (NS-398) and COX-2 (SC-560). We found that each inhibitor decreases fungal burden of H99-GFP infected larvae but not *Δplb1-*GFP infected larvae. Due to the phospholipase B1 dependence of this phenotype we think that these inhibitors might be affecting eicosanoid production by the *C. neoformans* itself. The ability of broad COX inhibitors like aspirin/indomethacin to inhibit eicosanoid production by *C. neoformans* is controversial (17,28) however our study is the first to use such selective COX-1 and COX-2 inhibitors on *C. neoformans*. This experiment remained inconclusive as to whether host PGE_2_ synthesis promotes virulence so to block PGE_2_ in zebrafish larvae without potential off target effects we used CRISPR Cas9 technology to knockdown expression of the prostaglandin E_2_ synthase gene *ptges* in zebrafish larvae. Ablating zebrafish *ptges* with this approach did not affect the fungal burden of *Δplb1-*GFP infected larvae but it did cause increased burden in H99-GFP infected zebrafish. This phenotype is interesting because it suggests host PGE_2_ might actually be inhibitory to cryptococcal virulence, furthermore this phenotype was phospholipase B1 dependent which suggests host-derived PGE_2_ might interact in some way with *Cryptococcus*-derived eicosanoids. This phenotype was not seen *in vitro* with aspirin so it is possible that the inhibitory effects of host-derived PGE_2_ influence a non-macrophage cell type in zebrafish larvae.

To confirm our observations that *C. neoformans* was the source of PGE_2_ during infection we performed co-infection assays with H99 wild type cryptococci (eicosanoid producing) and *Δplb1* (eicosanoid deficient) within the same macrophage and found that co-infection was sufficient to promote the intracellular growth of *Δplb1*. We also observed similar interactions during *Δlac1* co-infection (a second eicosanoid deficient *C. neoformans* mutant). These observations agree with previous studies that suggest eicosanoids are virulence factors produced by *C. neoformans* during macrophage infection (19,28). To identify if the secreted factor produced by *C. neoformans* was PGE_2_, we measured the levels of PGE_2_ from *Cryptococcus* infected macrophages to see if there was an observable increase in this eicosanoid during infection. Although PGE_2_ was detected, we did not see any significant difference between infected and uninfected macrophages, an observation confirmed using two different detection techniques – ELISA and LC MS/MS. These data suggest that PGE_2_ produced by *C. neoformans* during macrophage infection is contained within the macrophage, likely in close proximity to the fungus and that the host receptor targeted by these eicosanoids is therefore likely to be intracellular. Our *in vitro* co-infection experiments indicate that *C. neoformans* secretes virulence enhancing eicosanoids during infection.

The biological activity of 15-keto-PGE_2_ is far less studied than PGE_2_. It is known that 15-keto-PGE_2_ cannot bind to prostaglandin E_2_ EP receptors, this means it can act as a negative regulator of PGE_2_ activity i.e. cells up-regulate 15PGDH activity to lower PGE_2_ levels (47). Our findings however suggested that 15-keto-PGE_2_ did have a biological activity independent of PGE_2_ synthesis, possibly via a distinct eicosanoid receptor. It has been demonstrated that 15-keto-PGE_2_ can activate the intracellular eicosanoid receptor peroxisome proliferator associated receptor gamma (PPAR-γ) (32). Activation of PPAR-*γ* by *C. neoformans* has not been described previously but it is compatible with what we know of cryptococcal pathogenesis. PPAR-*γ* is a nuclear receptor normally found within the cytosol. Upon ligand binding PPAR-*γ* forms a heterodimer with Retinoid X receptor (RXR) and translocates to the nucleus where it influences the expression of target genes which possess a peroxisome proliferation hormone response element (PPRE) (48). If eicosanoids are produced by *C. neoformans* during intracellular infection, it is likely that they bind to an intracellular eicosanoid receptor. Additionally, activation of PPAR-*γ* within macrophages is known to promote the expression of anti-inflammatory genes which could make the macrophage more amenable to parasitism by the fungus.

To investigate whether PPAR-*γ* activation within macrophages occurs during *C. neoformans* infection we performed immunofluorescent staining of H99 and *Δplb1* infected J774 macrophages. We found that infection with *C. neoformans* led to increased nuclear localization of PPAR-*γ* indicating that the fungus was activating endogenous PPAR-*γ* during infection. We also found that macrophages infected with H99 had higher levels of PPAR-*γ* activation than *Δplb1* infected macrophages. This strongly suggests that eicosanoids produced by *C. neoformans* are responsible for activating PPAR-*γ*.

To confirm that 15-keto-PGE_2_ is an agonist to PPAR-*γ* we performed experiments with zebrafish larvae from PPAR-*γ* GFP reporter fish (36) and demonstrated that 15-keto-PGE_2_ binds to zebrafish PPAR-*γ*. To determine if 15-keto-PGE_2_ is an agonist to PPAR-*γ* we found that treating *Δplb1-*GFP infected zebrafish larvae with GW9662 at the same time as 15-keto-PGE_2_ blocked the virulence enhancing effects of the eicosanoid. These data indicate that 15-keto-PGE_2_ is a partial agonist to PPAR-*γ* (in zebrafish at least). Partial agonists are weak agonists that bind to and activate receptors, but not at the same efficacy as a full agonist. Partial agonists to PPAR-*γ* have been reported previously, partial PPAR agonists bind to the ligand binding domain of PPAR-*γ* with a lower affinity than full PPAR agonists and as a result activate smaller subsets of PPAR-*γ* controlled genes (38,39,49-52).

We also found that activation of PPAR-*γ* alone was sufficient to mediate cryptococcal virulence. In this respect, we found that the *in vitro* intracellular proliferation of the wild type H99 cryptococcal strain within J774 macrophages could be suppressed using a PPAR-*γ* antagonist GW9662. We could also block the rescuing effect of 15-keto-PGE_2_ on *Δplb1-*GFP during zebrafish infection using GW9662. Finally we found that *Cryptococcus* infected zebrafish treated with troglitazone at a concentration that is known to activate PPAR-*γ* (53) had increased fungal burdens when infected with *Δplb1-GFP*, *Δlac1-*GFP and H99-GFP strains. Taken together these experiments provide convincing evidence that a novel cryptococcal virulence factor - 15-keto-PGE_2_ – enhances the virulence of *C. neoformans* by activation of host PPAR-*γ* and that macrophages are one of the key targets of this eicosanoid during infection.

In this study, we have shown for the first time that eicosanoids produced by *C. neoformans* can promote virulence in an *in vivo* host. Furthermore, we have provided evidence that this virulence occurs via eicosanoid mediated manipulation of host macrophages. We have identified that the eicosanoid responsible for these effects is 15-keto-PGE_2_ which is derived from the dehydrogenation of PGE_2_ produced by *C. neoformans*. We have subsequently demonstrated that 15-keto-PGE_2_ mediates its effects via activation of PPAR-*γ*, an intracellular eicosanoid receptor known to promote anti-inflammatory immune pathways within macrophages. We provide compelling evidence that eicosanoids produced by *C. neoformans* enhance virulence, identifies a novel virulence factor – 15-keto-PGE_2_ – and describes a novel mechanism of host manipulation by *C. neoformans* - activation of PPAR-*γ*. Most importantly this study provides a potential new therapeutic pathway for treatment of cryptococcal infection, as several eicosanoid modulating drugs are approved for patient treatment (54).

## Materials and methods

(all reagents are from Sigma-Aldrich, UK unless otherwise stated)

### Ethics statement

Animal work was performed following UK law: Animal (Scientific Procedures) Act 1986, under Project License PPL 40/3574 and P1A4A7A5E. Ethical approval was granted by the University of Sheffield Local Ethical Review Panel. Experiments using the PPAR-γ reporter fish line (53) were conducted at the University of Toronto following approved animal protocols (# 00000698 “A live zebrafish-based screening system for human nuclear receptor ligand and cofactor discovery”) under a OMAFRA certificate.

### Zebrafish

The following zebrafish strains were used for this study: *Nacre* wild type strain, *Tg(mpeg1:mCherryCAAX)sh378)* transgenic strain and the double mutant *casper*, for PPARγ reporter experiments (53), which lacks all melanophores and iridophores (55). Zebrafish were maintained according to standard protocols. Adult fish were maintained on a 14:10 – hour light / dark cycle at 28 °C in UK Home Office approved facilities in the Bateson Centre aquaria at the University of Sheffield.

### C. neoformans

The H99-GFP strain has been previously described (31). The *Δplb1*-GFP and *Δlac1*-GFP stains was generated for this study by transforming existing deletion mutant strains (23,56) with a GFP expression construct (see below for transformation protocol). All strains used are in the *C. neoformans* variety *grubii* H99 genetic background.

*Cryptococcus* strains were grown for 18 hours at 28 °C, rotating horizontally at 20 rpm. *Cryptococcus* cultures were pelleted at 3300g for 1 minute, washed twice with PBS (Oxoidm Basingstoke, UK) and re-suspended in 1ml PBS. Washed cells were then counted with a haemocytometer and used as described below.

### *C. neoformans* transformation

*C. neoformans* strains *Δplb1* and *Δlac1* were biolistically transformed using the pAG32_GFP transformation construct as previously described for H99-GFP (31). Stable transformants were identified by passaging positive GFP fluorescent colonies for at least 3 passages on YPD agar supplemented with 250 μg/ml Hygromycin B.

### Zebrafish CRISPR

CRISPR generation was performed as previously described (29). Briefly gRNA spanning the ATG start codon of zebrafish *ptges* or *tyr* was injected along with Cas9 protein and tracrRNA into zebrafish embryos at the single cell stage. Crispant larvae were infected with *C. neoformans* as described above at 2 dpf. The genotype of each larvae was confirmed post assay – genomic DNA was extracted from each larvae and the ATG was PCR amplified with primers spanning the ATG site of *ptges* (Forward primer gccaagtataatgaggaatggg, Reverse primer aatgtttggattaaacgcgact) producing a 345-bp product. This product was digest with Mwol – wildtype digests produced bands at 184, 109 and 52 bp while mutant digests produced bands at 293 and 52 bp (Supplementary Figure 4).

### J774 Macrophage infection – with exogenous PGE_2_ treatment

J774 macrophage (J774 cells were obtained from the ATCC, American Type Culture Collection) infection was performed as previously described (21) with the following alterations. J774 murine macrophage-like cells were cultured for a minimum of 4 passages in T75 tissue culture flasks at 37°C 5% CO_2_ in DMEM (High glucose, Sigma) supplemented with 10% Fetal Bovine Calf Serum (Invitrogen), 1% 10,000 units Penicillin / 10 mg streptomycin and 1 % 200 mM L – glutamine, fully confluent cells were used for each experiment. Macrophages were counted by haemocytometer and diluted to a concentration of 1×10^5^ cells per ml in DMEM supplemented with 1 μg/ml lipopolysaccharide (LPS from *E. coli*, Sigma L2630) before being plated into 24 well microplates (Greiner) and incubated for 24 hours (37 °C 5% CO_2_).

Following 24-hour incubation, medium was removed and replaced with 1 ml DMEM supplemented with 2 nM prostaglandin E_2_ (CAY14010, 1mg/ml stock in 100% ethanol). Macrophage wells were then infected with 100 μl 1×10^6^ yeast/ml *Cryptococcus* cells (from overnight culture, washed. See above) opsonized with anti-capsular IgG monoclonal antibody (18b7, a kind gift from Arturo Casadevall). Cells were incubated for 2 hours (37 °C 5% CO_2_) and then washed with 37 °C PBS until extracellular yeast were removed. After washing, infected cells were treated with 1ml DMEM supplemented with PGE_2_.

To calculate IPR, replicate wells for each treatment/strain were counted at 0 and 18 hours. Each well was washed once with 1ml 37 °C PBS prior to counting to remove any *Cryptococcus* cells released by macrophage death or vomocytosis. Intra-macrophage Cryptococci were released by lysis with 200 μl dH_2_O for 20 minutes (lysis confirmed under microscope). Lysate was removed to a clean microcentrifuge tube and an additional 200 μl was used to wash the well to make a total lysate volume of 400 μl. *Cryptoccoccus* cells within lysates were counted by haemocytometer. IPR was calculated by dividing the total number of counted yeast at 18hr by the total at 0hr.

To assess the viability of *C. neoformans* cells recovered from macrophages we used our previously published colony forming unit (CFU) viability assay (21). Lysates from *C. neoformans* infected J774 cells were prepared from cells at 0hr and 18hr time points. The concentration of *C. neoformans* cells in the lysate was calculated by haemocytomter counting, the lysates were then diluted to give an expected concentration of 2×10^3^ yeast cells per ml. 100 μl of this diluted lysate was spread onto a YPD agar plate and incubated for 48 hr at 25°C prior to colony counting.

### J774 Macrophage co-infection

J774 cells were prepared and seeded at a concentration of 1×10^5^ per ml as above in 24 well microplates and incubated for 24 hours (37 °C 5% CO_2_), 45 minutes prior to infection J774 cells were activated with 150 ng/ml phorbol 12-myristate 13-acetate in DMSO added to 1 ml serum free DMEM. Following activation J774 cells were washed and infected with 100 μl / 1×10^6^ yeast/ml 50:50 mix of *Δplb1* (non-fluorescent) and H99-GFP (e.g. 5×10^5^ *Δplb1* and 5×10^5^ H99-GFP) or *Δlac1-GFP* and H99 (non-fluorescent). Infected cells were incubated for 2 hours (37 °C 5% CO_2_) to allow for phagocytosis of *Cryptococcus* and then washed multiple times with 37 °C PBS to remove unphagocytosed yeast, each well was observed between washes to ensure that macrophages were not being washed away. After washing 1 ml DMEM was added to each well.

Co-infected cells were imaged over 20 hours using a Nikon TE2000 microscope fitted with a climate controlled incubation chamber (37 °C 5% CO_2_) using a Digital Sight DS-QiMC camera and a Plan APO Ph1 20x objective lens (Nikon). GFP and bright field images were captured every 4 minutes for 20 hours. Co-infection movies were scored manually. For example co-infected macrophages that contained 2 *Δplb1* (non-fluorescent) and 1 H99-GFP (GFP positive) yeast cells at 0 hr were tracked for 18 hours and before the burden of each strain within the macrophage was counted again. The IPR for *Δplb1* within co-infected macrophages was calculated by dividing the number of *Δplb1* cells within a macrophage at 18 hr by the number at 0 hr.

### Immunofluorescence

J774 cells were cultured to confluency as discussed above and seeded onto sterile 13 mm glass coverslips at a density of 10^5^ cells per ml without activation by phorbol 12-myristate 13-*Δplb1*-GFP were opsonised with 18B7 for one hour. 18B7 was then removed by centrifugation and fungal cells were suspended in 1 ml of PBS with 1:200 FITC for one hour. Supernatant was removed again, cells were resuspended in PBS, and J774 were infected with 10^6^ of either H99-GFP or *Δpbl1*-GFP in serum free DMEM. After two hours of infection media was removed from J774s and the J774s were washed three times with PBS. 10μm TLT or DMSO was added to uninfected cells to act as controls. Cells were then left for 18 hours at 37 °C 5 % CO_2_.

After 18 hours supernatants were removed and J774s were fixed with cold methanol for 5 minutes at −20 °C before washing with PBS three times, leaving PBS for five minutes at room temperature between washes. Coverslips were blocked with 5% sheep serum in 0.1% triton (block solution) for 20 minutes before being transferred into the primary PPAR-γ antibody (1:50, γ Santa Cruz Biotechnology sc-7273 lot #B1417) with 1:10 human IgG in block solution for one hour. Coverslips were washed 3 times in PBS and incubated with 1:200 anti-mouse TRITC, 1:40 anti-human IgG, and 0.41 μl/ml DAPI in block solution for one hour. Coverslips were then washed three times with PBS, three times with water, and fixed to slides using MOWIOL. Slides were left in the dark overnight and imaged the following day. Imaging was performed on a Nikon Eclispe Ti microscope with a x60 DIC objective. Cells were imaged with filter sets for Cy3 (PPAR-γ, 500ms exposure) GFP (*Cryptococcus*, 35 ms exposure) and DAPI (Nuclei, 5ms) γ dyes in addition to DIC.

The intensity of nuclear staining was analysed for at least 30 cells per coverslip, using ImageJ 2.0.0 a line ROI was drawn from the outside of cell, through the nucleus measuring the mean grey value along the line. For *Cryptococcus* infected conditions uninfected and infected cells were measured separately upon the same coverslip using the GFP channel to distinguish between infected and uninfected cells.

### J774 aspirin timelapse

Macrophages were seeded at 10^5^ per ml into 24 well plates as described above. After two hours cells requiring aspirin were treated with 1 mM aspirin in DMSO in fresh DMEM. Cells were then incubated overnight for 18 hours at 37 °C 5% CO_2_. H99-GFP and *Δplb1*-GFP were prepared at 10^6^ cells per ml as described above, and opsonised with 18B7 for one hour. J774s were then infected with the fungal cells in fresh serum free DMEM for two hours before removing the supernatant, washing three times in PBS, and adding fresh serum free DMEM. Cells were imaged for 18 hours on a Nikon Eclispe Ti equipped with a climate controlled stage (Temperature - 37 °C, Atmosphere - 5% CO_2_ / 95% air) with a x20 Lambda Apo NA 0.75 phase contrast objective brightfield images were taken at an interval of 2 minutes, 50 ms exposure. Analysis was performed by manual counts of intracellular and extracellular cryptococci.

### Zebrafish infection

Washed and counted *Cryptococcus* cells from overnight culture were pelleted at 3300g for 1 minute and re-suspended in 10% Polyvinylpyrrolidinone (PVP), 0.5% Phenol Red in PBS to give the required inoculum in 1 nl. This injection fluid was loaded into glass capillaries shaped with a needle puller for microinjection. Zebrafish larvae were injected at 2-days post fertilisation; the embryos were anesthetised by immersion in 0.168 mg/ml tricaine in E3 before being transferred onto microscope slides coated with 3% methyl cellulose in E3 for injection. Prepared larvae were injected with two 0.5 nl boluses of injection fluid by compressed air into the yolk sac circulation valley. Following injection, larvae were removed from the glass slide and transferred to E3 to recover from anaesthetic and then transferred to fresh E3 to remove residual methyl cellulose. Successfully infected larvae (displaying systemic infection throughout the body and no visible signs of damage resulting from injection) were sorted using a fluorescent stereomicroscope. Infected larvae were maintained at 28 °C.

### Eicosanoid / receptor agonist treatment of infected zebrafish larvae

All compounds were purchased from Cayman Chemical. Compounds were resuspended in DMSO and stored at −20 °C until used. Prostaglandin E_2_ (CAY14010, 10mg/ml stock), Prostaglandin D_2_ (CAY12010, 10mg/ml stock), 16,16-dimethyl-PGE_2_ (CAY14750, 10mg/ml stock), 15-keto-PGE_2_ (CAY14720, 10mg/ml stock), troglitazone (CAY14720, 10mg/ml stock), GW9662 (CAY70785,1mg/ml stock), AH6809 (CAY14050, 1mg/ml stock), GW627368X (CAY10009162, 10mg/ml stock), NS-398 (CAY70590, 9.4mg/ml stock), SC-560 (CAY70340, 5.3mg/ml stock).

Treatment with exogenous compounds during larval infected was performed by adding compounds (or equivalent solvent) to fish water (E3) to achieve the desired concentration. Fish were immersed in compound supplemented E3 throughout the experiment from the time of injection.

### Zebrafish fungal burden measurement

Individual infected zebrafish embryos were placed into single wells of a 96 well plate (VWR) with 200 ul or E3 (unsupplemented E3, or E3 supplemented with eicosanoids / drugs depending on the assay). Infected embryos were imaged at 0 days post infection (dpi), 1 dpi, 2 dpi and 3 dpi in their 96 well plates using a Nikon Ti-E with a CFI Plan Achromat UW 2X N.A 0.06 objective lens. Images were captured with a Neo sCMOS (Andor, Belfast, UK) and NIS Elements (Nikon, Richmond, UK). Images were exported from NIS Elements into Image J FIJI as monochrome tif files. Images were threshholded in FIJI using the ‘moments’ threshold preset and converted to binary images to remove all pixels in the image that did not correspond to the intensity of the fluorescently tagged *C. neoformans*. The outline of the embryo was traced using the ‘polygon’ ROI tool, avoiding autofluorescence from the yolk sac. The total number of pixels in the threshholded image were counted using the FIJI ‘analyse particles’ function, the ‘total area’ measurement from the ‘summary’ readout was used for the total number of GFP^+^ pixels in each embryo.

### PPAR-*γ* GFP reporter fish treatment

PPARγ embryos were collected from homozygous ligand trap fish (F18). Embryos were raised in a temperature-controlled water system under LD cycle at 28.5° C. in 0.5×E2 media in petri-dishes till 1 dpf. Any developmentally delayed (dead or unfertilized) embryos were removed. Chorions were removed enzymatically with Pronase (1mg/ml) and specimens were dispensed into 24 well plates (10 per well) in 0.5×E2 media. For embryos 1% DMSO was used as a vehicle control. Chemicals were stored in DMSO, diluted appropriately and added individually to 400 ul of 0.5×E2 media with 0.05 U/ml penicillin and 50 ng/ml streptomycin and vortexed intensively for 1 min. 0.5×E2 media was removed from all wells with embryos and the 400 ul chemical solutions were administered to different wells to embryos. For drug treatment embryos were pre-incubated for 1 hour with compounds and then heat induced (28→37° C.) for 30 min in a → water bath. Embryos were incubated at 28°C for 18 h and then monitored using a fluorescent dissection scope (SteREO Lumar.V12 Carl Zeiss) at 2 dpf. For analyzing GFP fluorescent pattern, embryos were anesthetized with Tricaine (Sigma, Cat.# A-5040) and mounted in 2 % methyl cellulose.

### Whole body macrophage counts

2 dpf transgenic zebrafish larvae which have fluorescently tagged macrophages due to an mCherry fluorescent protein driven by the macrophage specific gene marker mpeg1 (57) *Tg(mpeg1:mCherryCAAX)sh378)* (22) were treated with 10 μM PGE_2_, 10 μM 15-keto-PGE_2_ or an equivalent DMSO control for 2 days. Larvae were then anesthetized by immersion in 0.168 mg/ml tricaine in E3 and imaged using a Nikon Ti-E with a Nikon Plan APO 20x/ 0.75 DIC N2 objective lens, taking z stacks of the entire body with 15 μM z steps. Macrophage counts were made manually using ImageJ from maximum projections.

### Eicosanoid measurement (ELISA)

J774 macrophages were seeded into 24 well plates at a concentration of 1×10^5^ per well and incubated for 24 hours at 37 °C 5 % CO_2_. J774 cells were infected with *C. neoformans* as described above, at the same MOI 1:10 and incubated for 18 hours with 1ml serum free DMEM. At 18 hours post infection the supernatant was removed for ELISA analysis.

For ELISA analysis with aspirin treatment, wells requiring aspirin had supernatants removed and replaced with fresh DMEM containing 1mM aspirin in 1% DMSO. Cells were left for 24 hours total. At 24 hours all wells received fresh serum free media. Wells requiring aspirin for the duration received 1mM aspirin in DMSO. Aspirin treated cells requiring arachidonic acid were treated with 30 μg per ml arachidonic acid in ethanol. Control wells received the following: either 1% DMSO, 30 μg per ml arachidonic acid, or ethanol. Cells were again left at 37 °C 5 % CO_2_ for 18 hours.

Supernatants were then removed and frozen at −80 °C until use. Supernatants were analysed as per the PGE_2_ EIA ELISA kit instructions (Cayman Chemical).

### Eicosanoid measurement (Mass spectrometry)

J774 macrophages were seeded into T25 tissue culture flasks at a concentration of 1.3×10^6^ cells per flask and incubated for 24 hours at 37 °C 5 % CO_2_. J774 cells were infected with *C. neoformans* as described above, at the same MOI 1:10 and incubated for 18 hours with 2 ml serum free DMEM. At 18 hours post infection infected cells were scraped from the flask with a cell scraper into the existing supernatant and immediately snap frozen in ethanol / dry ice slurry. All samples were stored at −80 °C before analysis.

Lipids and lipid standards were purchased from Cayman Chemical (Ann Arbor, Michigan). Deuterated standard Prostaglandin E_2_-d_4_ (PGE_2_-d_4_), ≥98% deuterated form. HPLC grade solvents were from Thermo Fisher Scientific (Hemel Hempstead, Hertfordshire UK).

Lipids were extracted by adding a solvent mixture (1 mol/L acetic acid, isopropyl alcohol, hexane (2:20:30, v/v/v)) to the sample at a ratio of 2.5 –1 ml sample, vortexing, and then adding 2.5 ml of hexane (58). Where quantitation was required, 2 ng PGE_2_-d_4_, was added to samples before extraction, as internal standard. After vortexing and centrifugation, lipids were recovered in the upper hexane layer. The samples were then re-extracted by addition of an equal volume of hexane. The combined hexane layers were dried and analyzed for Prostaglandin E_2_ (PGE_2_) using LC-MS/MS as below.

Lipid extracts were separated by reverse-phase HPLC using a ZORBAX RRHD Eclipse Plus 95Å C18, 2.1 × 150 mm, 1.8 μm column (Agilent Technologies, Cheshire, UK), kept in a column oven maintained at 45°C. Lipids were eluted with a mobile phase consisting of A, water-B-acetic acid of 95:5:0.01 (vol/vol/vol), and B, acetonitrile-methanol-acetic acid of 80:15:0.01 (vol/vol/vol), in a gradient starting at 30% B. After 1 min that was ramped to 35% over 3 min, 67.5% over 8.5 min and to 100% over 5 min. This was subsequently maintained at 100% B for 3.5 min and then at 30% B for 1.5 min, with a flow rate of 0.5 ml/min. Products were monitored by LC/MS/MS in negative ion mode, on a 6500 Q-Trap (Sciex, Cheshire, United Kingdom) using parent-to-daughter transitions of *m/z* 351.2 → 271.2 (PGE_2_), and *m/z* 355.2 → 275.2 for PGE_2_-d_4_. ESI-MS/MS conditions were: TEM 475 °C, GS1 60, GS2 60, CUR 35, IS −4500 V, dwell time 75 s, DP −60 V, EP −10 V, CE −25 V and CXP at −10 V. PGE_2_ was quantified using standard curves generated by varying PGE_2_ with a fixed amount of PGE_2_-d_4_.

## Acknowledgments

We thank the Bateson Centre aquaria staff for their assistance with zebrafish husbandry and the Johnston, Renshaw and Elks labs for critical discussions. We also thank Arturo Casadevall (Johns Hopkins University, Maryland USA) for providing the 18B7 antibody.

## Supplementary Fig 1

**A** Quantification of the vialibility of *C. neoformans* retrieved from the phagosomes of J774 macrophages at 2 hr and 20 hr post infection. Prior to, and during infection J774 macrophages were treated with 2 mM PGE_2_ or the equivalent amount of solvent (Ethanol) *Cryptococcus* cells were counted with a hemocytometer / diluted and plated to give an expected number of 200 CFU - dead *Cryptococcus* cells are indistinguishable from live cells when counting with a hemocytometer however a lower than expected CFU count would indicate that there is a decrease in viability. Data is displayed as the fold change between the CFU count at 2 hpi and 20 hpi for each condition. A one-way ANOVA with Tukey post test was performed comparing all conditions. H99 ETOH vs. *Δplb1* ** p = 0.0052, H99 ETOH vs. *Δplb1* ETOH ** p = 0.0056, H99 ETOH vs. *Δplb1* 2 mM PGE_2_ * p = 0.029. **B i** Comparison of fungal burden between H99-GFP, *Δplb1-*GFP and *Δlac1-*GFP infected larvae (Data reproduced from Fig2 Ai, Di and Supplementary Fig 2 Bi for clarity) H99-GFP, *Δplb1-*GFP and *Δlac1-*GFP infected larvae imaged at 0, 1, 2 and 3 dpi. At least 50 larvae measured per time point from 3 biological repeats. Box and whiskers show median, 5^th^ percentile and 95^th^ percentile. Unpaired Mann-Whitney U tests used to compare the burden between each strain for every time point. **B ii** Table of Mann-Whitney U tests comparing burden for each strain between time points. **C** Whole body macrophage counts of zebrafish larvae treated at 2 dpf with 10 μM PGE_2_, 10 μM 15-keto-PGE_2_ or an equivalent DMSO control for 2 days. Box and whiskers show median, 5^th^ percentile and 95^th^ percentile. At least 12 larvae quantified per treatment group per biological repeat n = 4. Mann-Whitney U test used to treatments to DMSO control ** p = 0.0025.

## Supplementary Fig 2

**A** J774 cells co-infected with a 50:50 mix of *Δlac1-*GFP and H99. Quantification of IPR for *Δlac1*-GFP cells within *Δlac1*-GFP:H99 2:1 or 1:2 co-infected macrophages. At least 20 co-infected macrophages were analysed for each condition over 4 experimental repeats. Student’s T test performed to compare ratios − 2:1 vs 1:2 * p = 0.012. **B i** *Δlac1*-GFP infected larvae imaged at 0, 1, 2 and 3 dpi. Fungal burden measured by counting GFP positive pixels in each larvae. At least 78 larvae measured per time point across 3 biological repeats. Box and whiskers show median, 5^th^ percentile and 95^th^ percentile. Unpaired Mann-Whitney U tests used to compare the burden between each strain for every time point, for p values see (Supplementary Fig2 A + B). **B ii** Representative GFP images (representative = median value) of 2dpi *Δlac1-*GFP infected larvae, untreated at 0,1,2,3 dpi **C i** *Δlac1*-GFP Infected larvae treated with 10 μM prostaglandin E_2_ or equivalent solvent (DMSO) control. At least 70 larvae measured per treatment group across 4 biological repeats. Box and whiskers show median, 5^th^ percentile and 95^th^ percentile. Unpaired Mann-Whitney U test used to compare between treatments, DMSO vs. 10 μM PGE_2_ * p = 0.035. **D i** *Δlac1*-GFP Infected larvae treated with 10 μM 16,16-dimethyl prostaglandin E_2_ or equivalent solvent (DMSO) control. At least 75 larvae measured per treatment group across 4 biological repeats. Box and whiskers show median, 5^th^ percentile and 95^th^ percentile. Unpaired Mann-Whitney U test used to compare between treatments, DMSO vs. 10 μM 16,16-dimethyl prostaglandin E_2_ ns p = 0.062. **E i** *Δlac1*-GFP Infected larvae treated with 10 μM 15-keto-prostaglandin E_2_ or equivalent solvent (DMSO) control. At least 58 larvae measured per treatment group across 3 biological repeats. Unpaired Mann-Whitney U test used to compare between treatments DMSO vs. 10 μM 15-keto-prostaglandin E_2_ ns p= 0.50. **C ii, D ii, E ii** Representative GFP images (representative = median value) *Δlac1-*GFP infected larvae, at 2 dpi treated with 10 μM PGE_2_ (**C ii**), 16,16-dm-PGE_2_ (**D ii**) or 15-keto-PGE_2_ (**E ii**).

## Supplementary Fig 3

**A** PGE_2_ monoclonal EIA ELISA performed on supernatants from *C. neoformans* infected macrophages collected at 18 hr post infection. Mean concentration of PGE_2_ (pg per 1×10^6^ cells) plotted with SD, n = 2**. B** Quantification of IPR for *Δplb1* cells within *Δplb1*:H99-GFP co-infected macrophages at initial burdens of 2:1, 3:1, 4:1 and vice versa. N=4. Student’s T test performed to compare ratios – 2:1 vs 1:2 * p = 0.0137. **C** Example images of immunofluorescence experiments in J774 macrophages staining for PPAR-gamma nuclear localization (for quantification see Fig 5 Ai). J774 cells treated with DMSO, 0.25 μM Troglitazone, infected with H99-GFP or *Δplb1-*GFP at x60 magnification, scale bar = 10 μM. Images provided are from the Cy3 channel (PPAR-γ) and the corresponding cell in DIC. The area of the nuclei is marked with a white dotted line.

## Supplementary Fig 4

Genotyping to confirm zebrafish *ptges* CRISPR. Zebrafish were genotyped post assay (5 dpf), an area of genomic DNA spanning the *ptges* gene ATG site (the CRISPR target) was amplified with PCR to produce a 345 bp product. This product was digested with Mwol to produce genotype specific banding patterns **A.** Schematic of the banding patterns expected for each genotype following Mwol digestion – 1. Undigested product, a single 345 bp band 2. Wild type genotype, 184, 109 and 52 bp bands 3. Hetrozygous genotype (*ptges* ^+/−^) 293, 184, 109 and 52 bp bands 4. Homozygous genotype (*ptges ^−/−^*) strong bands for 293 and 52 bp, weaker bands at 184 and 109 bp can sometimes be seen indicating a small amount of wildtype *ptges* is still present (this is thought to be beneficial as low levels of *ptges* are required for larvae survival. **B** Genotyping for H99 infected larvae, L = DNA ladder (NEB 50 bp ladder), wt = undigested wild type control, wt dig = wild type digested control, numbers correspond to individual larvae genotyped. H99 + tyro = *tyr*^−/−^ larvae infected with H99-GFP. H99 + ptges = *ptges*^−/−^ larvae infected with H99-GFP **C** Genotyping for *Δplb1* infected larvae, L = DNA ladder (NEB 50 bp ladder), wt = undigested wild type control, wt dig = wild type digested control, numbers correspond to individual larvae genotyped. *Δplb1*+ tyro = *tyr*^−/−^ larvae infected with *Δplb1*-GFP. *Δplb1*+ ptges = *ptges*^−/−^ larvae infected with *Δplb1*-GFP.

