## Supplementary figures and images for "15-keto-prostaglandin E_2_ activates host peroxisome proliferator-activated receptor gamma (PPAR-γ) to promote *Cryptococcus neoformans* growth during infection"

### Sup Fig 1

A

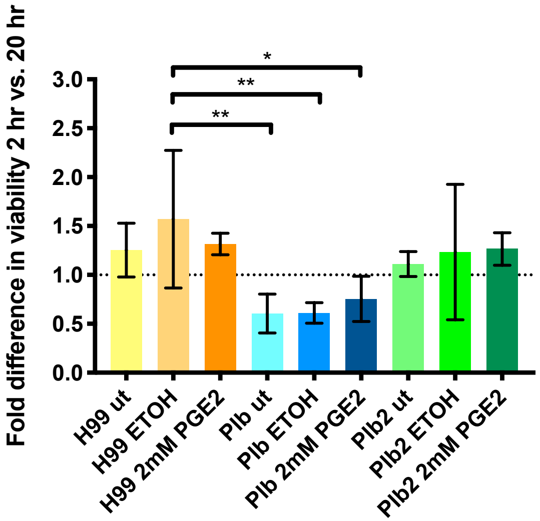


B i

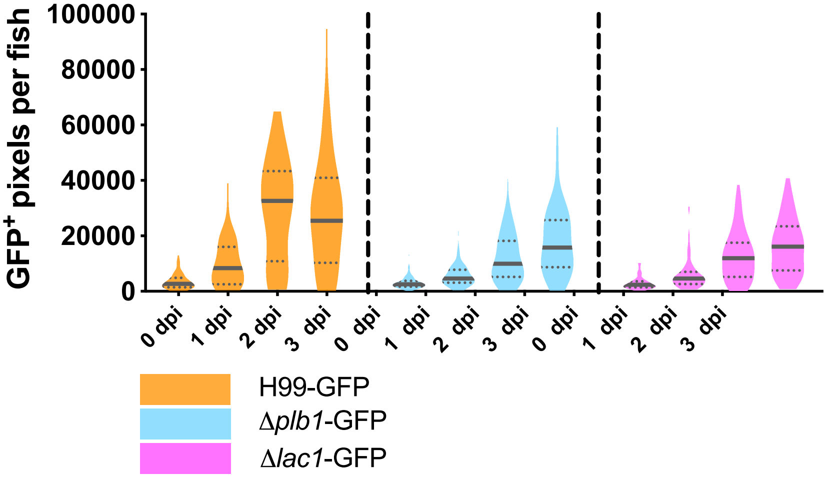


ii

| DPI (H99-GFP vs. *Δplb1*-GFP) | P value |
|---|---|
| 0 | ns p = 0.4980 |
| 1 | ** p = 0.0095 |
| 2 | **** p = <0.0001 |
| 3 | ** p = 0.0013 |
| **DPI (H99-GFP vs. *Δlac1*-GFP)** | **P value** |
| 0 | ns p = 0.2442 |
| 1 | ** p = 0.0088 |
| 2 | **** p = <0.0001 |
| 3 | *** p = 0.0002 |

C

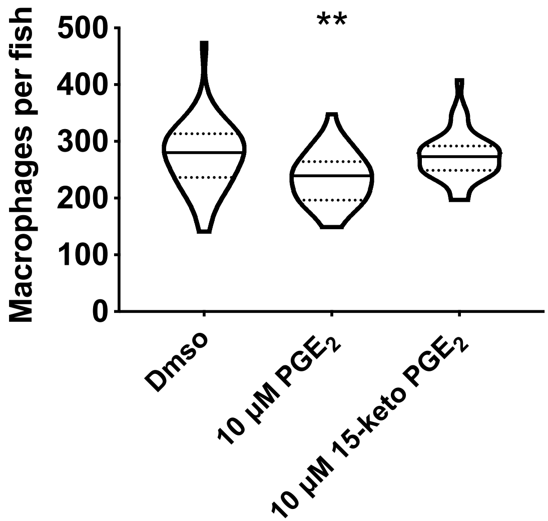

### Sup Fig 2

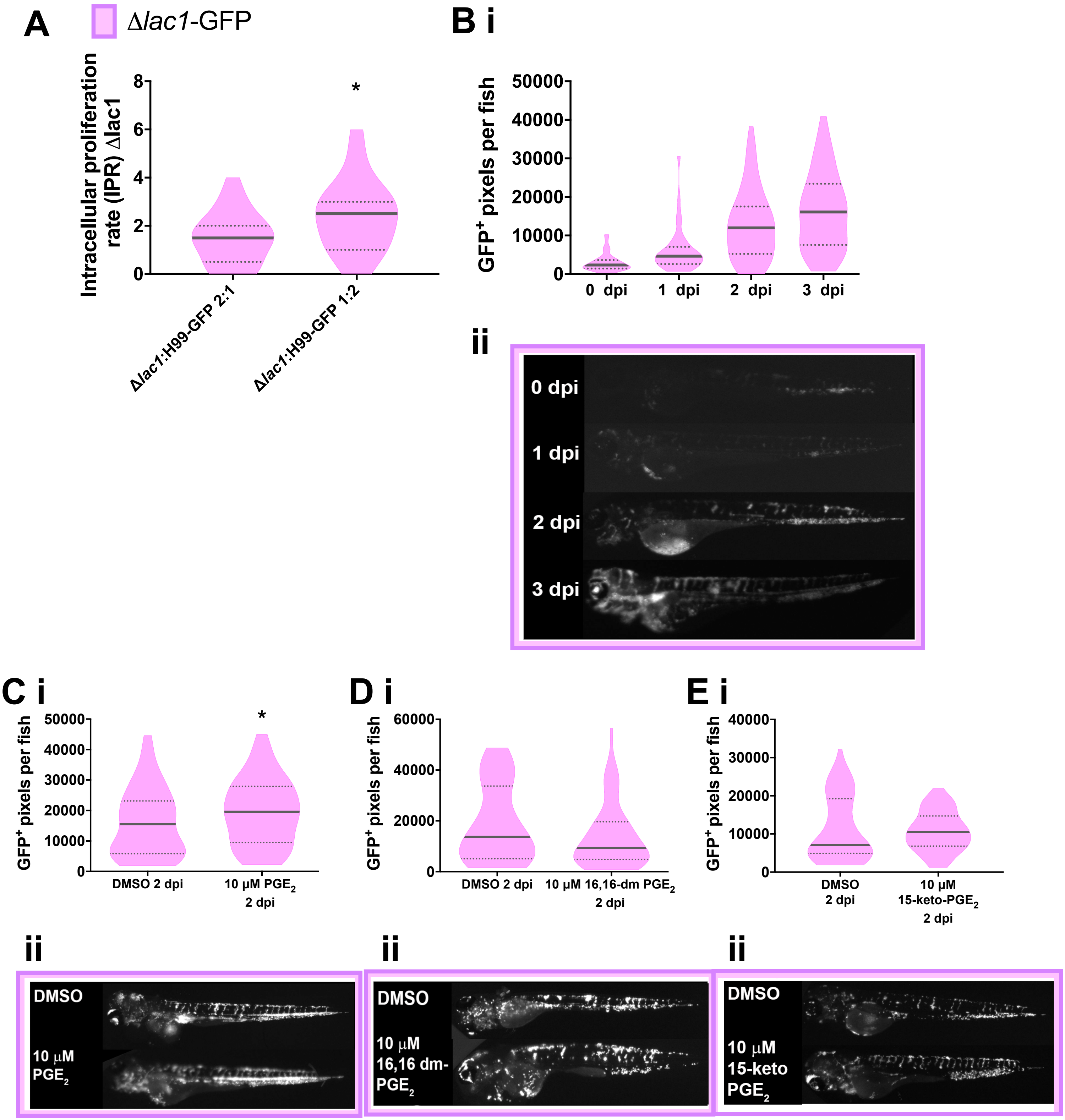

### Sup Fig 3

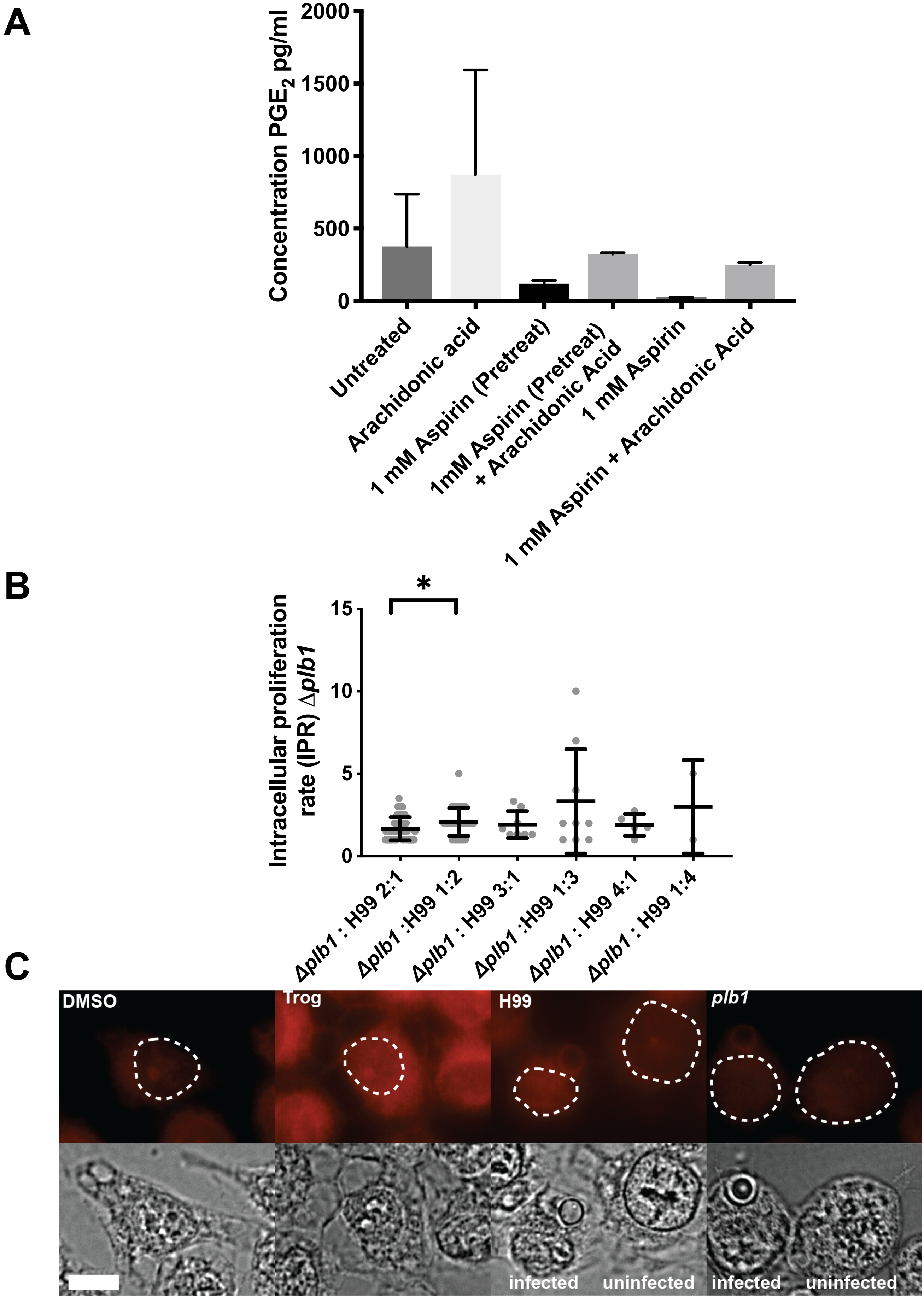

### Sup Fig 4

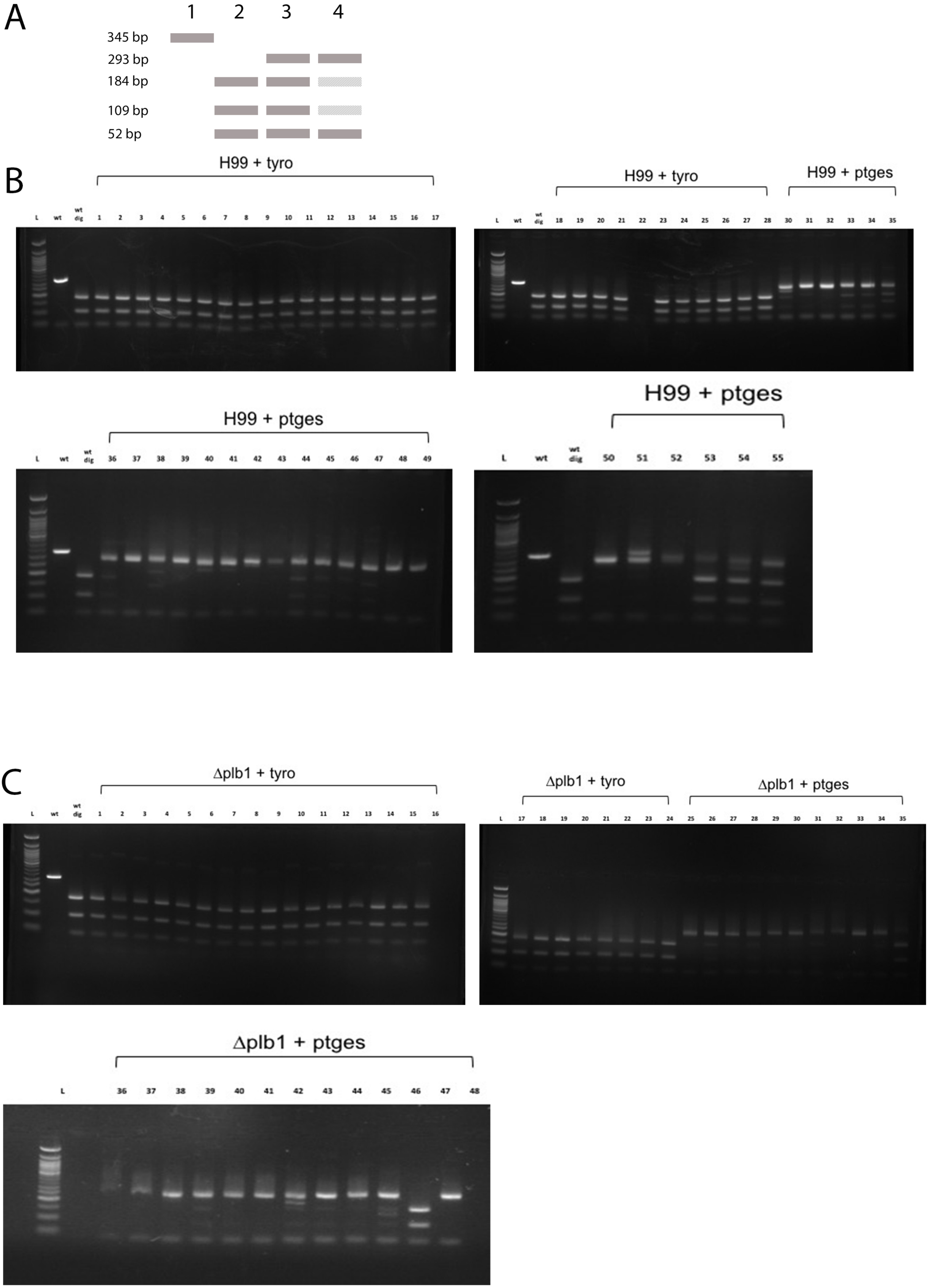
